# Presynaptic Terminals Dynamically Modulate Spontaneous Release Frequency During Early Synaptic Plasticity Through and Entropic Force Framework

**DOI:** 10.64898/2026.07.03.736394

**Authors:** Paxton Wilson, Henry Stephens, Rachel Cotter, Mira Menon, Brennan Plank, Miranda N. Reed, Michael W. Gramlich

**Author notes:** Authors contributed equally to this work (Paxton Wilson, Henry Stephens).

## Abstract

Spontaneous synaptic transmission has been established as essential for the maintenance of synaptic weights during action potential-induced transmission. However, spontaneous transmission also changes during synaptic plasticity and has been shown to, in part, mediate changes in synaptic weights. Despite decades of research, a coherent framework for understanding the complex molecular processes that support presynaptic spontaneous transmission during maintenance and plasticity has remained elusive. We show here that presynapses modulate spontaneous transmission frequency during the early time-course of plasticity following entropic force theory. We use live primary hippocampal cultures as a model system and induce plasticity using an established Long-Term Potentiation (LTP) protocol. We then use a combination of electron microscopy, fluorescence microscopy, and computational modeling to show how spontaneous release frequency dynamically changes during early plasticity. We use our entropic force theory to show how the dynamically changing synaptic vesicle pool structure mediates spontaneous release changes. Lastly, we show how these changes are altered in the presence of P301L tau leading to degeneration. The results from this study provide new insights that not only help understand normal synaptic function but also aid in understanding neurodegeneration.

## Introduction

Synapses represent the fundamental unit of information transfer in the central nervous system. They accomplish this task at the molecular level via an action potential-mediated release of neurotransmitter (NT) from a presynapse into the synaptic cleft, where the NT binds to postsynaptic receptors, initiating an ionic current (which we will call stimulated release).^1–5^ The complex molecular interactions that underlie this process are governed by a well-established, experimentally supported multinomial model.^6–8^ This multinomial model has also served as a basis for understanding changes in neuronal function under normal and neurodegenerative conditions, ^9,10^ as well as for underpinning computational neuroscience and artificial intelligence algorithms in the form of synaptic weights.^10–12^

A major limitation with the multinomial model is the so-called stability/plasticity balance, which is the competition between maintaining (stability) and modifying (plasticity) synaptic weights.^13,14^ This problem has been difficult to solve because of the complex combination of molecular-level changes that occur both in the presynapse^15–17^ and postsynapse^18–21^ between stimulated release events. Thus, over the last couple of decades, the role of changing synaptic behavior between stimulated release events has become a major focus.

One long-hypothesized pathway supporting synaptic maintenance has been spontaneous synaptic transmission.^22–24^ In the absence of an action potential, presynapses will spontaneously release NT at a regular frequency.^23,25–27^ Despite the fact that spontaneous release uses similar molecular machinery (such as synaptic vesicles and release sites), no coherent model framework has been developed to explain or predict spontaneous release frequency. Developing a model framework has been difficult because changes in spontaneous release mechanics have been shown to be unrelated to changes in stimulated release.^22^ We recently showed that an entropic force model framework can predict spontaneous release frequency.^28^ This framework is based on the theory that the energy density of the NT carrying synaptic vesicles (SVs) exerts force on the SVs tethered at the membrane, driving spontaneous fusion events similar to an ideal gas exerting pressure on the walls of a container. Further, using this framework, we showed that amyloid-beta mutations alter spontaneous release frequency during homeostasis.

Interestingly, spontaneous NT release has also been shown to mediate synaptic plasticity.^22,28–30^ For example, long-term potentiation (LTP) protocols have been shown to induce a short-term, significant increase in spontaneous release frequency, followed by a slow reduction back toward baseline; however, the post-LTP induction release frequency remains above baseline.^31^ Separately, SV distribution and density have also been shown to change >30 min after LTP induction.^32,33^ While these recent studies have used high-resolution electron microscopy, few have examined the early time-course of dynamic changes, which may not follow a simple trend and can have long-reaching implications. This gap in understanding the time-course of plasticity is essential to overcome to fully understand how plasticity occurs. Further, a single coherent framework that can predict how presynaptic spontaneous release mediates plasticity, or vice versa, has yet to be distinguished. Establishing a time-dependent predictive framework is essential because deficits in both spontaneous release and plasticity have been shown to correlate with neurodegeneration.^34–36^

In this study, we distinguish how SV spontaneous release frequency changes during early stages of presynaptic plasticity using our entropic force model. We use cultured primary hippocampal mouse neurons as a model system. We first induce synaptic plasticity using an established theta-burst protocol shown to correlate with associative memory,^37^ changes in SV distribution,^32,33^ and spontaneous release.^31^ We use electron microscopy to show that SV distribution and density change dynamically over time and with synapse size. Based on SEM results, we predict changes in spontaneous release frequency using the entropic force framework and. We then use an experimental pHluorin-VGlut1 approach to support entropic force theory predictions that show spontaneous release frequency dynamically changes, consistent with a change in SV density. We use a single SV SGC5 labeling approach to show that SV mobility is constrained within the presynapse, and the dynamically changing SV number is regulated via inter-synaptic vesicle exchange (ISVE). We last show how these changes are altered in the presence of P301L tau, known to lead to frontal temporal dementia. Thus, we observe a potential mechanism that mediates memory deficits and neurodegeneration prior to observable changes.

## Results

### Section 1: Presynapse SV Distribution Changes During Early Plasticity

Presynaptic spontaneous release rate depends on the density of vesicles near the active zone under homeostatic conditions.^38^ However, previously published studies have also shown spontaneous vesicle release and SV distribution within the presynapse change after plasticity induction.^21,33,39^ To establish any potential changes during early plasticity and test whether the entropic force theory can relate changes in SV density to spontaneous release and predict spontaneous release rates, we first sought to experimentally determine how SV number and density change early after plasticity induction. We used co-cultured primary hippocampal astrocytes and neurons (**Supplementary Methods**, **Fig. 1A**), given their essential role in memory formation and maintenance.^40^ We induced plasticity using the well-established *θ*-burst protocol that is utilized in long-term potentiation (LTP) (**Supplementary Methods, Fig. 1B**).^33,37,41,42^ In this study, we will refer to this protocol and the time-dependent presynaptic changes as induction of plasticity (or simply ***induction***). We then waited a fixed amount of time (<20 min, which we will call **LTP**) after induction to fix, stain, slice, and/or image samples using our established Large Area SEM (La-SEM) protocol (**Supplementary Methods, Fig. 1C**).^28,43^

**Figure 1:**
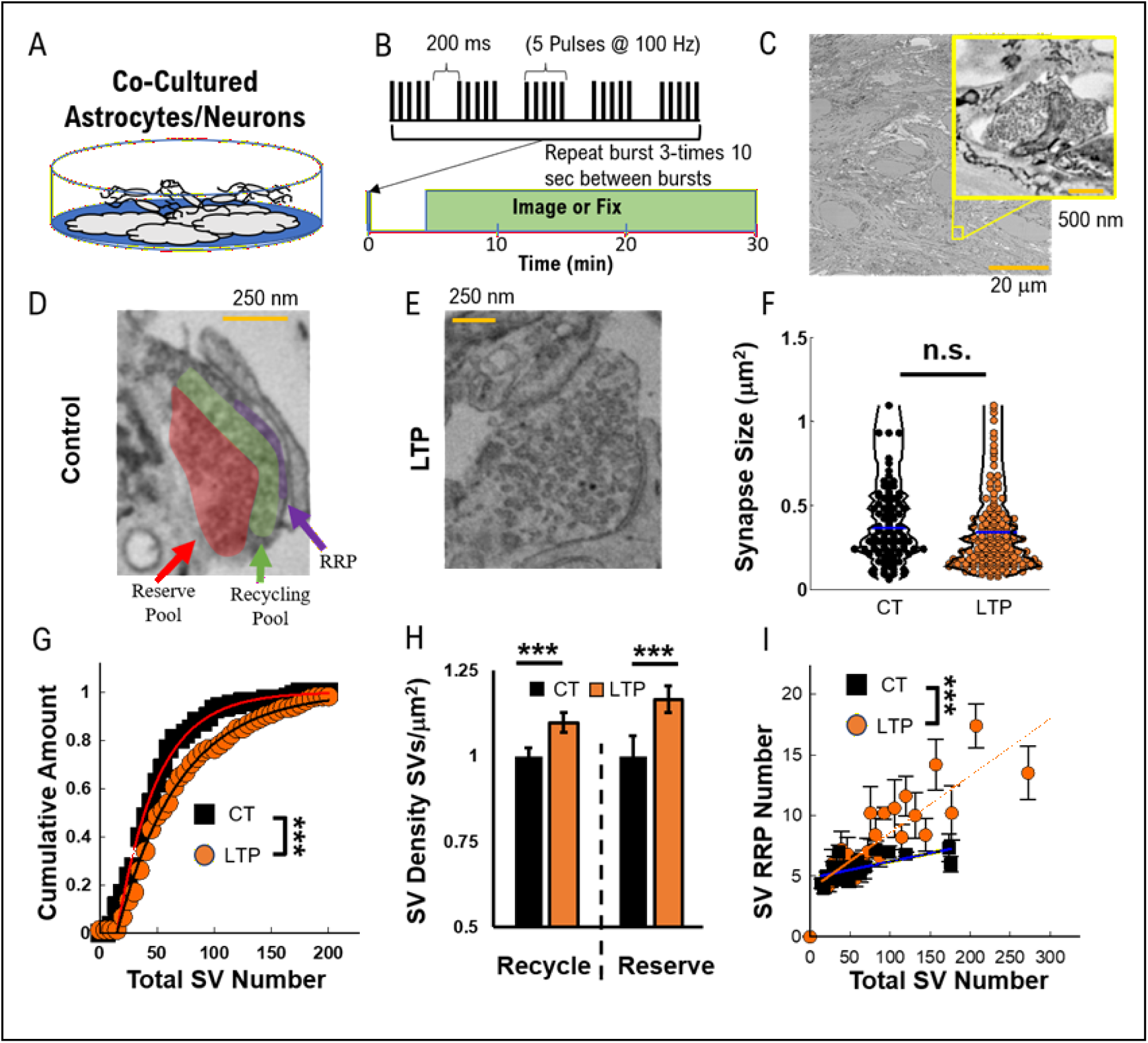
Cultured hippocampal neurons exhibit changing SV number and density during plasticity. (A) We utilized co-cultured hippocampal neurons/astrocytes to quantify synaptic function. (B) We induced potentiation-based plasticity using the established *θ*-burst used in Long-Term Potentiation (LTP) protocols. (**C**) We fixed, stained, sliced, and imaged cultures using established SEM approaches. We observe a changing SV number from baseline (**Control, D**) in the RRP (Purple region) Recycling Pool (Green Region) and Reserve Pool (Red Region), increasing at <20 min after induction (**LTP, E**). (**F**) Overall presynapse size did not significantly change after LTP induction (Orange) compared to baseline (Black). (**G**) The total number of SVs doubled during LTP (Orange Circles) above baseline (Black Squares). (**H**) We then separated presynapses into recycling and reserve groups and observed an increase in relative SV density both near the plasma membrane (Recycling) and farther away (Reserve) after induction (**LTP, Orange**). (**I**) We further compared the number of SVs at the membrane (RRP) and observed a linear increase after induction (**LTP, Orange Circles**) as a function of total SV number. Control data from two cultures with synapses N_ct_ = 111; LTP data from two cultures with total synapses N_LTP_ = 165 Statistical comparisons are: (G) a two-tailed KS-test; (H)(I) a two-tailed t-test for all experimental data. * = P< 0.05; ** = P < 0.01; *** = P< 0.001. Data points represent: (F) & (H) average of all synapses, and (I) moving average of SV number over 5 synapses.

We observed a qualitative increase in the number of SVs per presynapse after induction (**Fig. 1D-E**). To determine the synapse size changes, we quantified the average synapse size and found no significant overall change (**Fig. 1F**) and a slight average decrease. We then quantified the change in SV number and observed a significant increase in SV number relative to baseline (**Fig. 1G**). It is essential to highlight that comparing the total SV number to synapse size shows that the same geometric size synapses exhibit more SVs during LTP (**Supplementary Fig. S1A**). Further, the rate of increase in the number of SVs as a function of presynapse size is greater after induction (Lines, **Supplementary Fig. S1A**). These combined results suggest that presynaptic terminals retain their geometric size even as the number of SVs increases. Interestingly, to our knowledge, this dynamic change in the total number of SVs has not previously been reported at this timescale.

While this SV number change has not previously been observed, a relative increase in SVs in the recycling pool has been observed in hippocampal slices 30 min after induction.^32,33^ To determine whether a dynamically changing SV number supports the previous observation, we quantified SV density as a function of distance from the AZ using our previous approach.^28^ We observed that, across all presynapses, SV density in both the recycling and reserve pools significantly increased after induction (**Fig. 1H**). Further, the fraction of SVs in the ready releasable pool increases after induction for the same number of total SVs per synapse compared to control (**Fig. 1I**). These results show that both the number and density of SVs within a presynapse significantly increase in the early time-course of plasticity.

### Section 2: Computational Entropic Force Model Predicts Increasing Spontaneous Release

To understand the consequences of a dynamically changing SV number/density on presynaptic function, we use the entropic force framework to estimate how spontaneous release rate changes in response to a changing SV distribution (**See Methods for derivation**). During homeostasis (**Fig. 2A**), spontaneous release frequency follows the relationship:

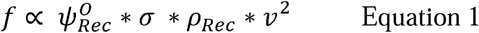

where *σ* is the number of release sites, *ψ* is the spontaneous translation parameter (SPOT), *ρ*_rec_ is the **density** of SVs in the recycling pool, v is the SV diffusive speed.

**Figure 2:**
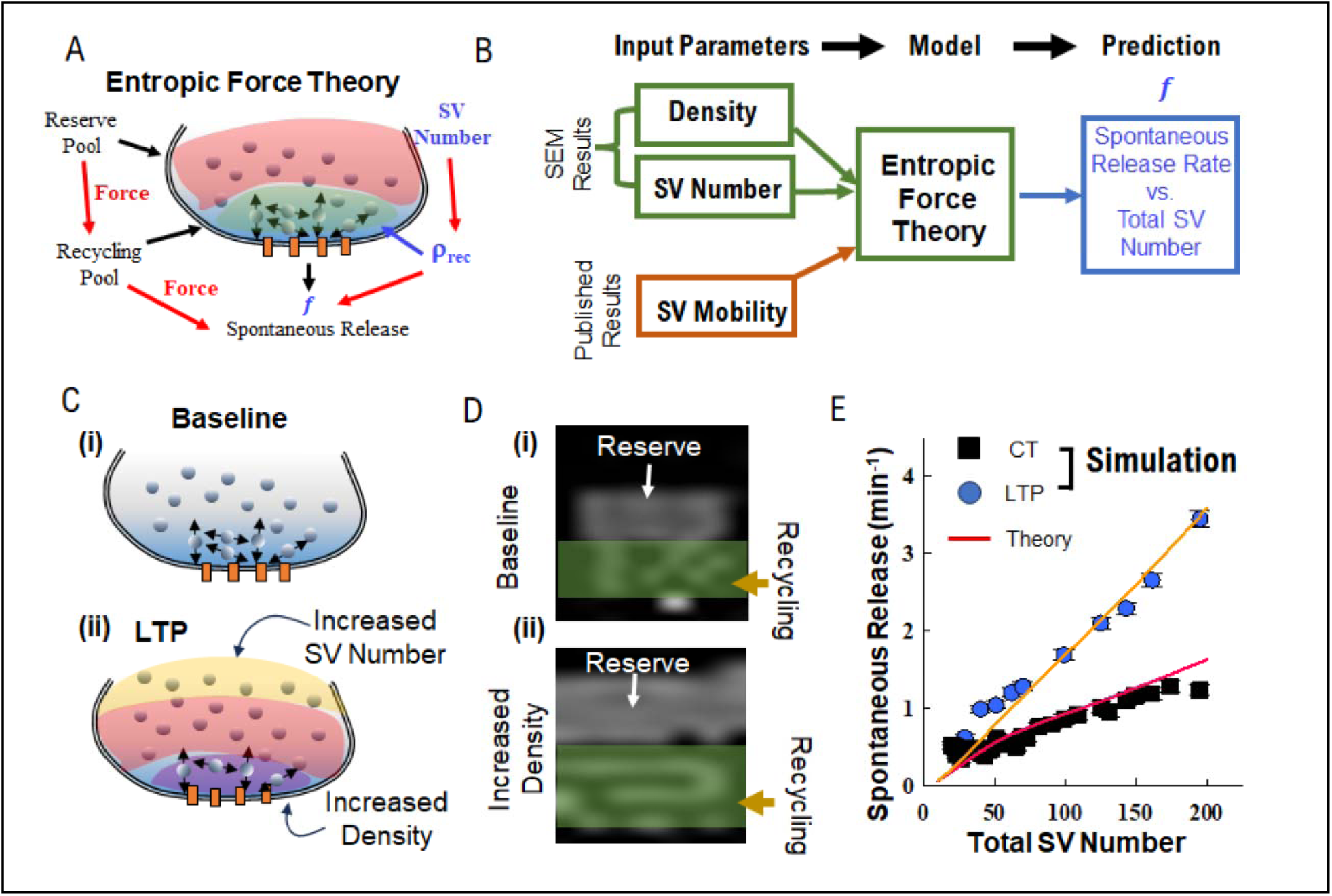
Entropic Force Theory Predictions from SEM Results. (A) Entropic Force theory proposes that SV density (*ρ*_rec_) is the mediating factor that leads to spontaneous release rate (Equation 1). (B) Entropic Force theory predicts spontaneous release rate (f) based on input parameters from SEM results and previously published SV recycling pool speed. (C) We hypothesize that (i) baseline SV density in the recycling pool increases during LTP (ii), leading to entropic force increasing the spontaneous release rate. (D) We computationally modeled changing SV density and mobility following a computational model of entropic force theory. (i) We first modeled the average SV density and mobility as a function of SV number at baseline, and (ii) then modeled the hypothesized increase in SV density during LTP. (E) Both computational simulations and entropic force theory (Solid lines) show that spontaneous release frequency increases 10 min after induction (Blue circles) compared to baseline (Black squares)

We used the SEM density and total SV number results (**Fig. 1G,H**) combined with previous experimental results,^28^ as input parameters for the model **(Fig. 2 B)** to predict:

i. The spontaneous release rate increases with total number of SVs per synapse
ii. If the observed presynaptic SV density dynamically changes during plasticity (**LTP, Fig. 1J**), then spontaneous release rate will change by a corresponding amount for the same size synapse.

We propose that, for presynapses of the same geometric size (**Fig. 2C(i)**), the increase in SV density during LTP will lead to an increase in spontaneous release rate (**Fig. 2C(ii)**). This is based on the experimental SEM observations (**Fig. 1F-I**) showing an increase in SV density for presynapses of the same size.

To show how the entropic force model can predict spontaneous release rates, we used a computational simulation of the model to estimate spontaneous release rates based on observed SV density changes in La-SEM data. We used the experimentally observed SV distribution measured using La-SEM (**Fig. 1G-I**, **Supplementary Fig. S1)**. We then put those distributions into the computational model, while holding the Spontaneous Translation (SPOT) parameter at homeostatic value. We then compared the resulting simulated spontaneous release rates for each observed state ( (i) Baseline, (ii) LTP, **Fig. 2D**).

The simulated presynapses exhibit an overall **linear** increase in the spontaneous release rate with increasing presynapse SV number (**Fig. 2E**). The model predicts that the spontaneous release rate after induction (Blue, **Fig. 2E**) is significantly higher than that of control synapses (Black, **Fig. 2E**). The spontaneous release rate follows our entropic force model (Red and Orange Solid lines, **Fig. 2E**). Further, it is essential to highlight that the LTP spontaneous release rate is predicted to exhibit a linear increase with SV number. These predictions suggest that recycling pool density near the active zone is a significant mediating factor in dynamically changing spontaneous release rates observed during plasticity.

These combined experimental and theoretical results *hypothesize* that a changing spontaneous release is mediated by a changing presynaptic SV *density*. For the remainder of this study, we will experimentally test the hypothesized spontaneous release rates and the mechanisms that mediate a dynamically changing SV distribution.

### Section 3: Spontaneous SV Exocytosis Rate Dynamically Changes During Plasticity

To determine if SV spontaneous exocytosis rate changes during plasticity as predicted by the entropic force model (**Fig. 2E**), we used our previously established pHluorin-VGlut1 protocol to measure spontaneous release.^28^ Briefly, we transduced hippocampal neurons with a pH-sensitive fluorescent reporter (**Fig. 3A**) and measured spontaneous release, low-frequency-stimulated release (1 Hz), high-frequency-stimulated release (40 Hz), and NH_4_Cl-induced release. This protocol allows us to quantify spontaneous release frequency, release probability, recycling pool SV number, and total SV number, respectively (**Fig. 3B**) as independent parameters within the same presynapse but on a synapse-by-synapse basis. We can then directly relate every aspect of synapse function to distinguish the mechanisms that mediate spontaneous release. We used the same induction protocol as in SEM (**Fig. 1B**).

**Figure 3:**
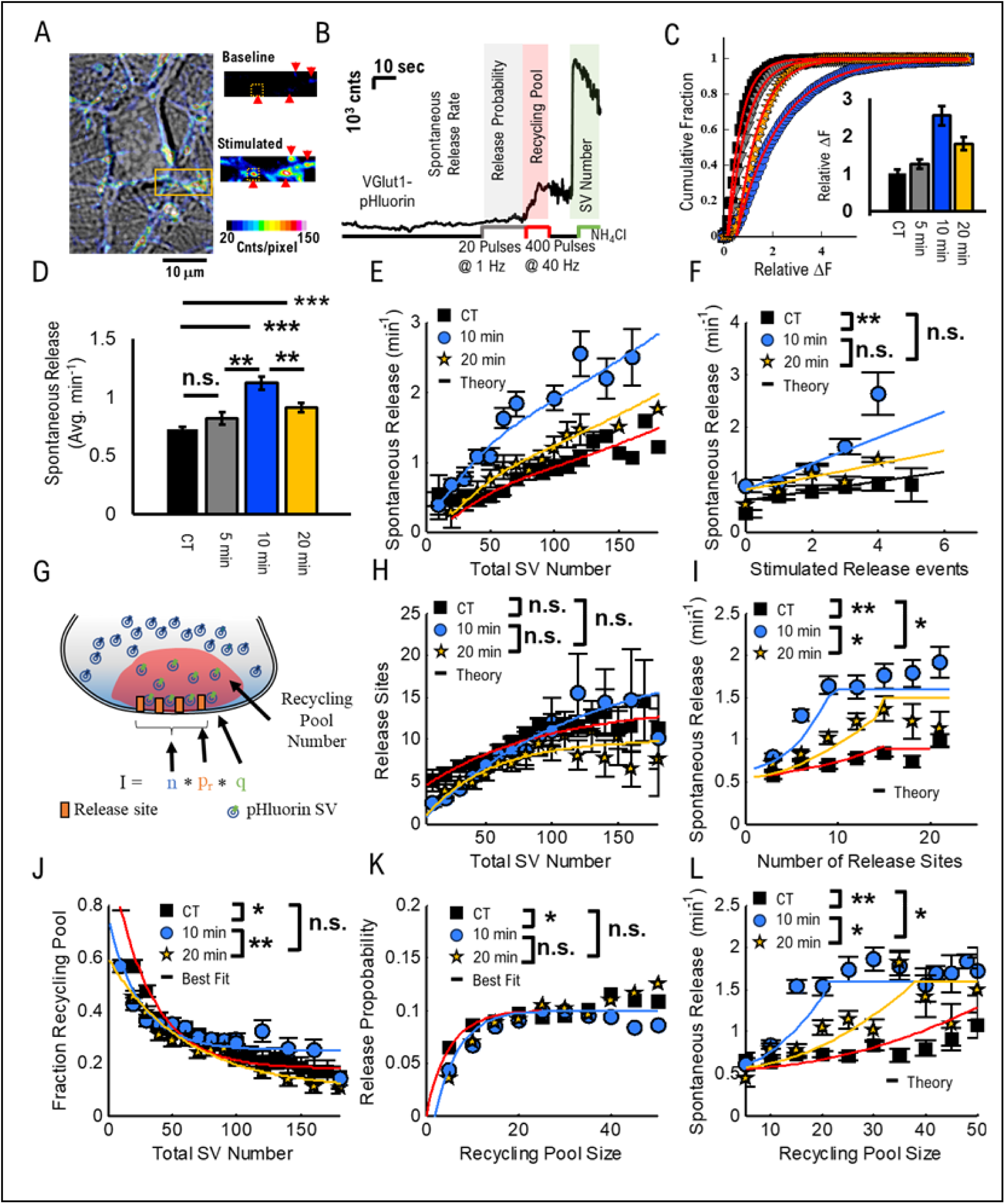
Presynaptic Spontaneous Release Rate Dynamically Changes During Plasticity. (**A**) Example cultured neuron viewed through transmitted light (Grey scale) and overlayed with pHluorin-VGlut1 intensity during stimulation (Color). Presynapses have low baseline fluorescence but exhibit higher fluorescence during stimulation due to fusing SVs, leading to changes in pH and fluorescence. The integration region around a single synapse (Dashed Yellow box) is used to quantify presynaptic SV intensity. (**B**) We quantify integrated intensity around each presynapse (Yellow Dashed Box, A) during a four-step stimulation protocol (See Methods). (**C**) We observe a significant and dynamic change in average presynaptic intensity as a function of time after induction. The pHluorin-VGlut1 intensity slightly increases at 5 min (Grey) and significantly at 10 min (Blue), followed by a decrease at 20 min (Yellow) toward baseline (Black). (**D**) We quantified the average spontaneous release rate across the network of synapses and observed a dynamic change as a function of time after induction. Spontaneous release slightly increases at 5 min (Grey) and significantly increases at 10 min (Blue), followed by a decrease at 20 min (Yellow) toward baseline (Black). (**E**) The spontaneous release rate as a function of total SV number shows a significant increase at 10 min after induction (Blue circles) followed by a reduction at 20 min after induction (Yellow Stars) compared to baseline (Black squares). Spontaneous release follows entropic force theory (Solid lines) at all time-points after induction. The entropic force theory dynamically adjusted the volume of the recycling pool while keeping all other parameters fixed based on electron microscopy results (Fig. 1). (**F**) Spontaneous release rate significantly increases as a function of stimulated release number for 10 min after induction (Blue Circles) and slightly 20 min after induction (Yellow Stars) compared to baseline (CT, Black Squares). All time points follow the entropic force model (Solid lines). (**G**) Cartoon multinomial release model used to estimate the number of release sites and recycling pool size from 40 Hz data (B) based on measured release probability (p_r_) and intensity per vesicle (q). (**H**) The rate of change in release site number as a function of total SV number increases linearly up to 100, then remains relatively constant thereafter for all time-points. (**I**) Spontaneous release rate significantly increases as a function of release site number for 10 min after induction (Blue Circles) and 20 min after induction (Yellow Stars) compared to baseline (CT, Black Squares). The 10 min data exhibit a maximum spontaneous release rate equivalent to a maximum SV packing density at 10 total release sites. All curves follow the entropic force model (Solid Lines). (**J**) The general relationship of the fraction of SVs in the recycling pool as a function of Total SV Number decreases exponentially (Solid Lines) and does not change significantly after induction. However, the distribution is shifted 10 min and 20 min after induction. (**K**) Release probability increases exponentially as a function of Recycling Pool Size up to 20 SVs, where it becomes constant (Solid lines). The overall Release Probability does not change significantly after plasticity induction. (**L**) Spontaneous release probability follows entropic force theory as a function of Recycling pool size (Solid Lines). Spontaneous Release plateaus at ∼1.5 min-1 at 10 min (Blue Circles) and 20 min (Yellow Stars) after induction due to the maximum packing density for the recycling pool observed in La-SEM (Fig. 1H). For panel (C), distribution of intensities were compared from samples taken at the same date from the same cultures and N > 1000 synapses for each condition. For panels (D)-(L) CT from 4 samples from 3 cultures: N = 2057. 5 min from 2 samples from 2 cultures: N = 336 10 min from 3 samples from 3 cultures: N = 642. 20 min from 2 samples from 2 cultures: N = 554. Statistics: (D)(E) 2-sampled t-test, (F)-(L) Mann-Whitney-U test. * = P< 0.05; ** = P < 0.01; *** = P< 0.001. Data points represent average binned values (See Methods). Error bars represent standard error of binned data average.

We first experimentally tested whether the total number of SVs changed dynamically over time after induction, as observed by SEM (**Fig. 1G**). We used the total *Δ*F increase in pHluorin-VGlut1 intensity during NH_4_Cl exposure (**Fig. 3C**) because it has been established as a measure of total SVs within a presynapse. We observed a slight increase at 5 min (Grey Triangles, **Fig. 3C**), followed by a significant increase at 10 min after induction (Blue Circles, **Fig. 3C**) compared to controls (Black squares, **Fig. 3C**). The intensity then began to decrease at 20 min (Orange Stars, **Fig. 3C**) toward control. We quantified the changing intensity (Solid Lines) and found that the intensity more than doubled at 10 min after induction (inset, **Fig. 3C**).

Entropic force theory predicts that a change in SV density (**Fig. 2E**) mediates a change in spontaneous release rather than a changing SV number alone. To test whether the pHluorin-VGlut1 measured change in SV number correlates with a change in spontaneous release frequency, we compared the overall average network-level spontaneous release frequency, following standard approaches. We observed that spontaneous release slightly increases 5 min after induction (Grey, **Fig. 3D**), peaks at 10 min (Blue, **Fig. 3D**), and returns toward baseline 20 min after induction (Orange, **Fig. 3D**). These results are both qualitatively and quantitatively consistent with the dynamic change in SV number observed with SEM (**Fig. 1G**) and NH4Cl (inset, **Fig. 3C**).

We then utilized a novel analysis of spontaneous release frequency, by quantifying the spontaneous release as a function of total number of SVs as the independent variable, on a synapse-by-synapse basis (**Supplementary Methods** and **Fig. 3E**). The importance of this approach is that it allows us to compare two independently measured parameters (spontaneous release and total SV number) in the same synapse, which the entropic force model hypothesizes are linearly dependent. If spontaneous release increases linearly with total SV number, then that supports the entropic force hypothesis. We observed the same linear increase in spontaneous release rate as a function of total SV number previously reported (Black squares, **Fig. 3E**). However, we also observed a significant scale shift higher in spontaneous release frequency at 10 min after induction (Blue Circles, **Fig. 3E**), for the same SV number. The spontaneous release frequency then decreased at 20 min after induction (Orange Stars, **Fig. 3E**) but remained above control levels. The scale shift increase is consistent with the simulated change from SEM results (Fig. 2E) hypothesized using the entropic force model. This result supports that the spontaneous release rate is mediated by the SV density proposed by the entropic force model.

We next addressed whether spontaneous release frequency is higher than that of action potential-induced SV release events, suggesting that spontaneous and action potential-induced release mechanisms are separate, as previously hypothesized during baseline.^22^ We compared the spontaneous release frequency as a function of the number of stimulated release events (**Fig. 3F**), because this measurement isolates changes in action potential-induced release from spontaneous release.^28^ Here, we observed that spontaneous release frequency increased significantly 10 min after induction (Blue Circles, **Fig. 3F**) compared to control (Black squares, **Fig. 3F**). The increased spontaneous release frequency began to decrease at 20 min after induction (Orange Stars, **Fig. 3F**), but remained above control. These results show that changes in spontaneous release mechanics during plasticity are distinct from those induced by action potentials.

### Section 4: Active Zone Mechanics and Recycling Pool Size Do Not Significantly Change During Early Stages of Plasticity

To determine how spontaneous release rates change dynamically during plasticity, we quantified the number of release sites and recycling-pool SVs (**Fig. 3G**) using our established multinomial model fit to the 40Hz stimulation intensity.^28,44,45^ Briefly, we used the single SV release intensity (q) and release probability (pr) based on the experimentally measured values during spontaneous release and 1 Hz stimulated release data, respectively. We then input those parameters in the multinomial model to fit the 40 Hz pHluorin-VGlut1 intensity while allowing the number of release sites and SVs in the recycling pool to vary as fit parameters. We then compared experimentally measured parameters (spontaneous release rate, stimulated release rate, and total SV number).

We next established if the number of release sites dynamically changed during plasticity and observed that there is a slight, but not statistically significant, decrease in the number of release sites at 10 min (Blue Circles, **Fig. 3H**) and 20 min after induction (Orange Stars, **Fig. 3H**), relative to control (Black squares, **Fig. 3H**). We hypothesize that this is because the number of SVs increases faster than the active zone release site formation. This may meaning that, for a given presynapse, the number of SVs increases during plasticity while the number of release sites remains constant, resulting in a lower measured release site number during pHluorin-VGlut1 (**Fig. 3H**).

We next measured the spontaneous release frequency as a function of release site number to determine if spontaneous release mechanics changed independent of release site mechanics. We observed a significant increase in spontaneous release frequency for both 10 min (Blue Circles, **Fig. 3I**) and 20 min after induction (Orange Stars, **Fig. 3I**), relative to control (Black squares, **Fig. 3I**). However, the spontaneous release frequency remained constant above 10 release sites for both 10 min and 20 min data (Solid Lines, **Fig. 3I**). Based on entropic force theory, this result suggests that the density of the recycling pool SVs remains constant or decreases at synapses with larger active zone sizes (Solid Lines, **Fig. 3I**).

To establish spontaneous release frequency changes relative to recycling pool number, we quantified the fraction of the recycling pool as a function of total SV number (**Fig. 3J**). We observed a slight decrease in the fraction of recycling pool SVs at 10 min (Blue Circles, **Fig. 3J**) and 20 min after induction (Orange Stars, **Fig. 3J**), relative to control (Black squares, **Fig. 3J**) for synapses with 50 SVs or smaller. This is consistent with the release site results and *suggests that synapses increase the total number of SVs faster than they can increase the recycling pool number*.

We observe the same relative change in stimulated release probability as a function of Recycling Pool Number (**Fig. 3K**), with a scale shift at smaller recycling pool size. The stimulated release probability decreased for synapses with ∼10 recycling pool SVs at 10 min (Blue Circles, **Fig. 3H**) and 20 min after induction (Orange Stars, **Fig. 3K**), relative to control (Black squares, **Fig. 3K**). However, synapses with larger recycling-pool SVs are unchanged, likely due to stable release probability (pr ∼ 0.1) regardless of synapse size (>20 SVs), as observed under control conditions (Black squares, **Fig. 3K**). These results show that a vast majority of synapses do not have a significant change in their release probability during the early stages of plasticity.

We last turned to quantifying spontaneous release frequency as a function of recycling pool SV number. As with every other metric, we observe a significant increase in spontaneous release frequency for both 10 min (Blue Circles, **Fig. 3L**) and 20 min after induction (Orange Stars, **Fig. 3L**), relative to control (Black squares, **Fig. 3L**). Further, spontaneous release frequency begins to decrease 20 min after baseline but remains higher than control. Interestingly, the spontaneous release frequency remains relatively constant (∼1.75) for synapses with more than 20 SVs in the recycling pool at 10 min. This result supports the entropic force hypothesis: the recycling pool number alone does not mediate spontaneous release frequency; rather, the density of recycling pool SVs is the important mediating factor.

These combined pHluorin results show that spontaneous release frequency is significantly increased during plasticity, as predicted by the entropic force model using the initial SEM results (**Fig. 2B,E**). Further, the increase in spontaneous release does not correlate with increases in stimulated release, active zone release sites, or the SV number of the recycling pool. This leaves the hypothesized density of the recycling pool SVs as the mediating factor for the increase in spontaneous release, which we address next.

### Section 5: Synaptic Vesicle Mobility Within the Presynapse Dynamically Changes During Plasticity

We next sought to determine if the volume of the recycling pool SVs dynamically changes during the same conditions that the SV number dynamically changes during plasticity. Entropic force theory hypothesizes that the density of the recycling pool mediates the increase in spontaneous release rate during plasticity (**Fig. 3 D,E**). We used the established single-SV SGC5 labeling approach in rat hippocampal cultures (**See methods**), which we have previously shown can quantify the volume of the recycling pool.^28^ Briefly, we followed the same *θ*-burst and delay protocol (**Fig. 1B**) followed by a single SV labeling and imaging protocol (**Fig. 4A, See Methods**). We were able to image single SVs within presynapses (**Fig. 4B**). We then used our previously established correlation analysis algorithm^25,46^ to distinguish different types of motion within single SV tracks (**Fig. 4C**). Finally, we used the mobile fraction of SVs to determine whether SV mobility supports the dynamically changing spontaneous release rate.

**Figure 4:**
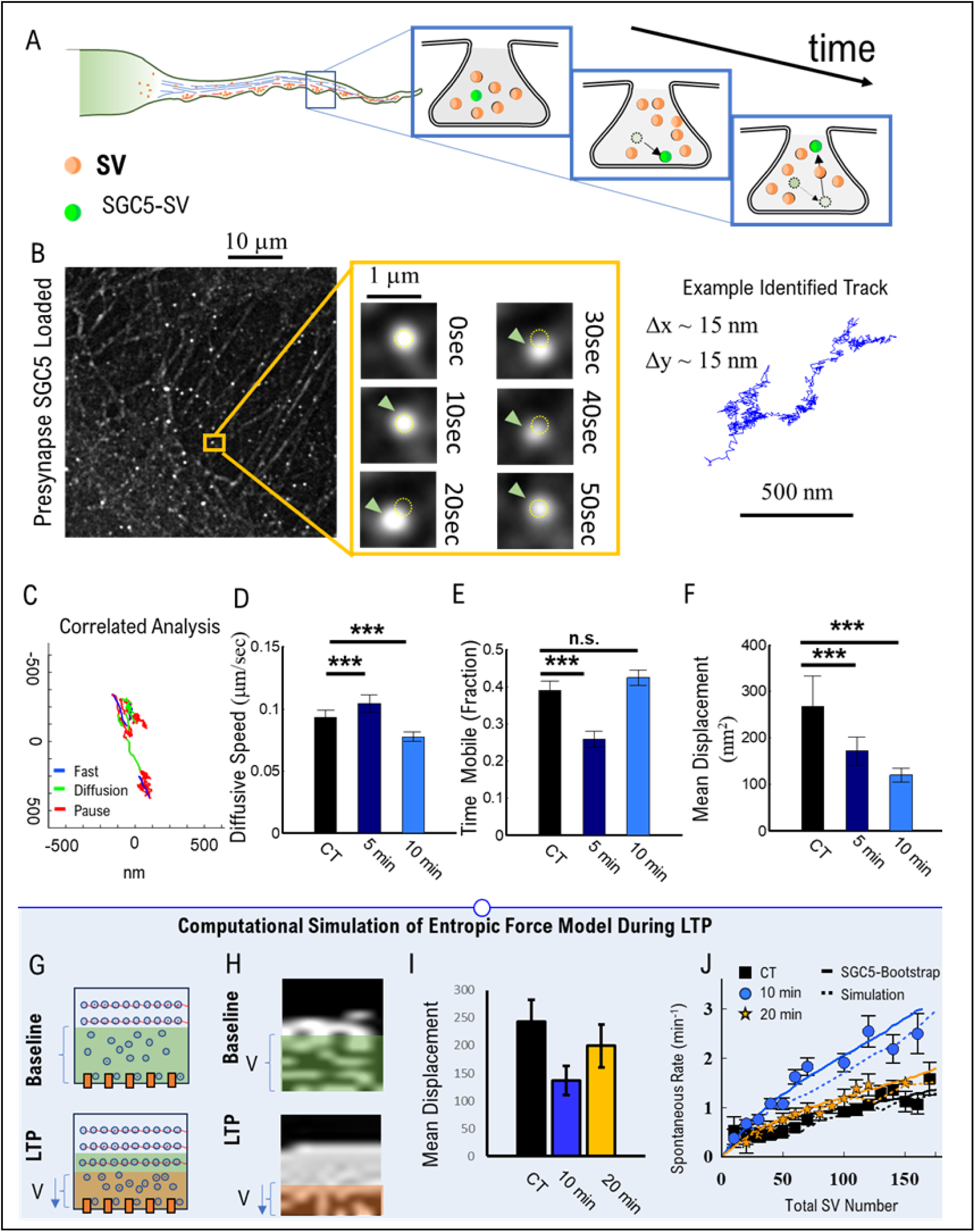
Single Labeled SVs mobility dynamically changes during plasticity to support changing spontaneous release rates. (A) Cartoon Depiction of labeled SGC5 SVs imaging approach within a presynapse. (**B**) Example frame of cultured hippocampal neurons labeled with SGC5 (Left) and an example of a single SGC5 labeled SV (middle) in hippocampal cultures. An example track (Right) from an identified SGC5 labeled SV with localization precision (*Δ*x,*Δ*y) to within 15 nm per frame. (**C**) An example track run through the correlation analysis algorithm identifying bouts of Fast, Diffusive, and Pause motion (See Methods). (**D**) The diffusive speed of SVs exhibits a slight increase at 5 min after induction (Dark Blue), followed by a slight decrease at 10 min (Light Blue). (**E**) The total time SVs are mobile decreases at 5 min after induction (Dark Blue), followed by a return to baseline at 10 min (Light Blue). (**F**) The mean displacement of high mobility SVs (See Methods) progressively decreases with time after plasticity induction. This result suggests that the volume of the recycling pool decreases with time after induction. (**G**) Cartoon model of presynaptic volume of the recycling pool dynamically changing during LTP (bottom) compared to baseline (top) based on mean displacement (F). (H) Snapshots of computational models of SV mobility within a presynapse under baseline (Top) and at 10 min after induction (Bottom). Spontaneous release events occur when SVs reach the bottom surface (See methods). (I) The simulated mean displacement of SVs away from the active zone as a function of time after plasticity. (J) Inferred spontaneous release rates from bootstrapped results of single SGC5 tracks (solid lines) and computational simulations (dashed lines) compared to pHluorin-VGlut1 measured spontaneous release rates for control (Black), 10 min (Blue Circles), and 20 min (Gold Stars) after baseline. CT: N_SVs_ = 104, from 7 samples; 5 min: N_SVs_ = 61, from 6 samples; 10 min: N_SVs_ = 196, from 6 samples; Statistics: (D)(E) 2-sampled t-test, (F) Mann-Whitney-U test. * = P< 0.05; ** = P < 0.01; *** = P< 0.001. Data points represent average values of SV. Error bars represent standard error of average.

We observed that SV mobility within presynapses changed after induction. First, SV diffusive speed decreased at 10 min after *θ*-burst (Blue, **Fig. 3D**), compared to control (Black, **Fig. 4D**). Alternatively, the fraction of time SVs spend mobile (Blue and Green color of track, **Fig. 4C**) decreased 5 min after the *θ*-burst (Blue, **Fig. 4E**), but returned to baseline at 10 min (Blue, **Fig. 4E**) compared to control (Black, **Fig. 4E**). Taken alone, these two results combined suggest that SVs in the recycling pool traverse the same total distance (distance = speed **x** time). However, speed and fraction of mobility do not indicate the displaced volume over which recycling pool SVs travel, which we have previously shown is the contributing factor in spontaneous release rate.^28^

To determine the volume over which the recycling pool SVs are constrained, we used the mean-square-displacement approach. Briefly, Recycling SVs are constrained in volume they traverse by the reserve pool structure,^47^ leading to a plateau in the MSD over time; ^28^ we then compare the maximum MSD to estimate the total volume of the recycling pool. We have previously shown that this metric decreased in the presence of Forskolin,^28^ a drug that is known to mimic LTP and synaptic plasticity. Here, we observe a halving in SV displacement 10 min after induction (Blue, **Fig. 4F**) compared to control (Black, **Fig. 4F**), which is consistent with our previous Forskolin observations.^28^ Further, this result is consistent with previously observed changes in recycling pool location in hippocampal slices after LTP.^33^ These results suggest that the recycling pool volume significantly reduces during plasticity, which the entropic force model predicts would lead to an increase in spontaneous release rate.

To establish whether the observed decrease in SV displacement (**Fig. 4F**) is sufficient to reproduce the observed increase in spontaneous release rate observed with pHluorin-VGlut1 (**Fig. 3E**), we used a combination of computational modelling and bootstrapping single SV experimental tracks (**Fig. 4C**) to predict the spontaneous release rate as a function of total SV number. We modeled SV mobility under a constrained volume, consistent with a 20-25% reduction in the recycling pool volume. This resulted in a slightly reduced instantaneous SV speed with a significant reduction in SV displacement (**Fig. 4I**). We then calculated the spontaneous release rate as a function of total SV number from the simulated tracks (blue and yellow dotted lines, **Fig. 4J**) and found that the reduction in recycling pool volume reproduced both the 10 min (blue circles, **Fig. 4J**) and 20 min (Yellow stars, **Fig. 4J**) pHluorin-VGlut1 results. We then used the SGC5 track data to reproduce the expected spontaneous release rate using entropic force theory and found that the tracks also reproduced the pHluorin-VGlut1 results (blue and yellow solid lines, **Fig. 4J**).

These combined results show that the increase in SV density observed with La-SEM (**Fig. 1H**) correlates with the reduction in recycling pool volume observed with SGC5 (**Fig. 4F**), which entropic force theory hypothesizes results in the increase in spontaneous release rate per SV with pHluorin-VGlut1 (**Fig. 3E**).

### Section 6: P301L Tau Neurons Do Not Exhibit Dynamical Changes During Plasticity

To determine whether experimentally observed dynamic changes during synaptic plasticity are altered during neurodegeneration, we used an established mouse model carrying the human P301L tau mutation.^48–52^ This model exhibits early-stage plasticity deficits and hyperexcitability at the network level. These changes have been proposed to correlate with increased VGLUT1 expression per synaptic vesicle (**Fig. 5B**).^52–55^ leading to increased glutamate release, which, in turn, causes hyperexcitability and seizure-like states.^56–61^ We previously showed that P301L neurons exhibit the same spontaneous and stimulated release, as well as synaptic function, as normal neurons under homeostatic conditions.^28^ However, *no previous study has explored if recycling pool changes contribute to plasticity deficits in P301L expressing neurons*.

**Figure 5:**
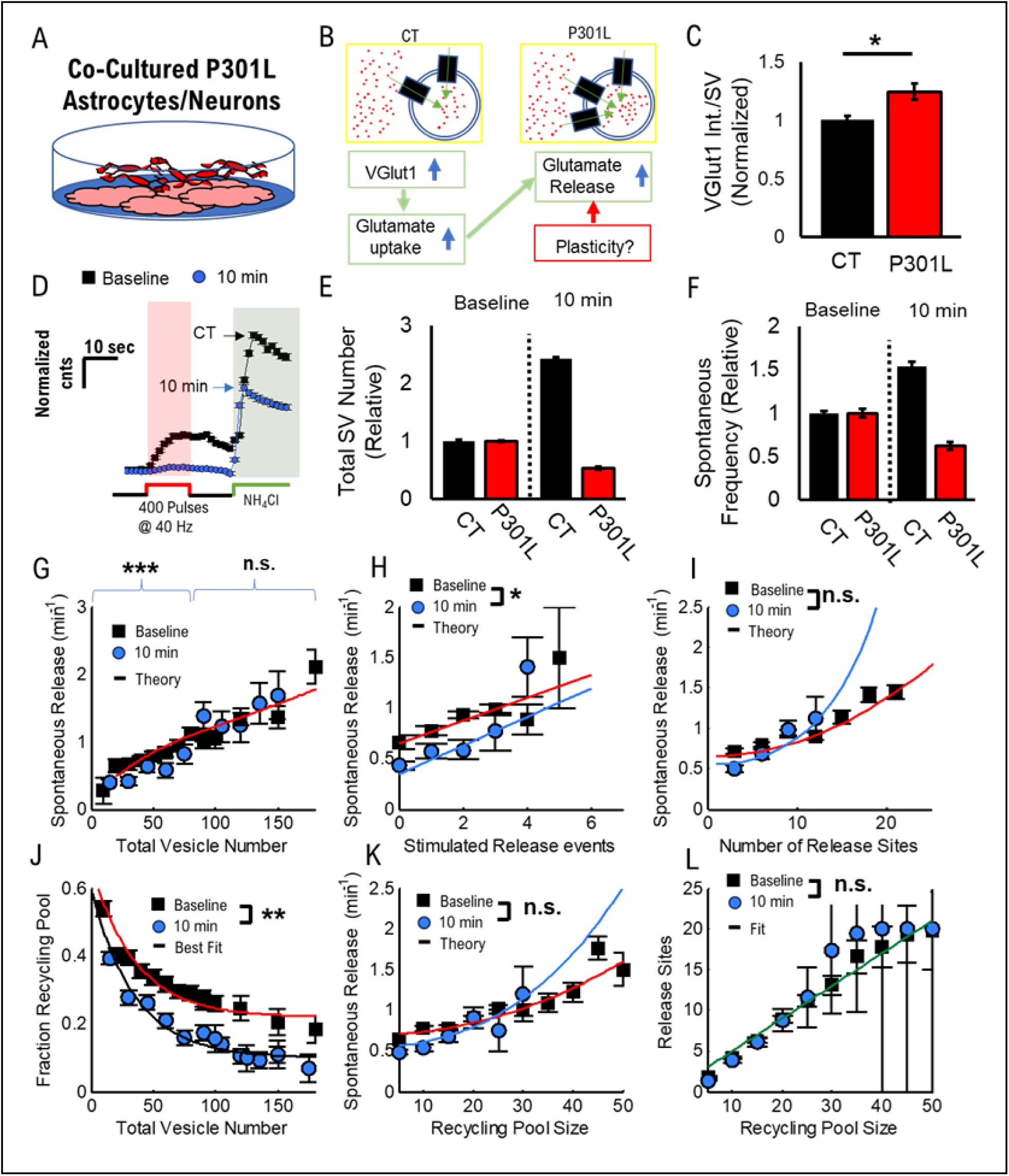
Spontaneous Release during plasticity is reduced in the presence of P301L due to loss of recycling pool SVs. (A) Cartoon of Co-cultured neurons/astrocytes. (B) Cartoon model of increased VGlut1 expression per vesicle. Established causal pathway of increased glutamate release and potential alterations during plasticity. (C) Comparison of VGlut1-pHluorin spontaneous release intensity for control (Black) and P301L (Red) cultures. The average P301L intensity is significantly increased, consistent with previous studies. (D) Average VGlut1-pHluorin intensity for P301L cultures under baseline (Black) and 10 min after plasticity induction (Blue Circles). P301L cultures exhibit a significant reduction in SV release during 40Hz stimulation and NH4Cl. (E) The estimated total number of SVs quantified from NH4Cl intensity distributions for control cultures (Black) and P301L cultures (Red). Total SV number increases 10 min after stimulation (Black, Right-Hand Side) compared to baseline (Black, Left-Hand Side) for control neurons. Total SV number decreases 10 min after stimulation (Red, Right-Hand Side) compared to baseline (Red, Left-Hand Side) for control neurons. (F) Average spontaneous release frequency for control cultures (Black) and P301L cultures (Red). Spontaneous release frequency significantly increases 10 min after stimulation (Black, Right-Hand Side) compared to baseline (Black, Left-Hand Side) for control neurons. Spontaneous release frequency significantly decreases 10 min after stimulation (Red, Right-Hand Side) compared to baseline (Red, Left-Hand Side) for control neurons. (G) Spontaneous release frequency per total SV is reduced at 10 min after stimulation (Blue Circle) for smaller presynapses (<70 SVs) compared to baseline (Black Squares) and unchanged for larger presynapses (>70 SVs). (H) Spontaneous release frequency per stimulated release is significantly reduced at 10 min after stimulation (Blue Circle) compared to baseline (Black Squares). (I) Spontaneous release frequency per release site is significantly reduced at 10 min after stimulation (Blue Circle) for smaller presynapses (<10 release sites) compared to baseline (Black Squares). (J) The fraction of SVs in the recycling pool is significantly reduced at 10 min after stimulation (Blue Circle) compared with baseline (Black Squares) across all presynaptic sizes. The best fit (solid lines) follows an exponential distribution as a function of SV number. (K) Spontaneous release frequency per recycling pool size is significantly reduced at 10 min after stimulation (Blue Circle) for smaller presynapses (<20 Recycling SVs) compared to baseline (Black Squares). Comparing the data to the entropic force model, the baseline data match the model (Red line) using the same volume parameters for normal neurons (Fig. 3L), while the 10 min data match the model (Blue line) using the same volume parameters from induced neurons (Yellow Line, Fig. 3L). (L) The number of release sites does not significantly change between baseline (Black squares) and 10 min after activity (Blue Circles). Sample Size: (C) CT means from 7 samples from 3 cultures; N>1000 SVs P301L means from 8 samples from 3 cultures; N>1000 SVs. *Note: Data imaged on the same date from cultures transducted on the same date and using the same microscope/media conditions. (E) - (F) CT Baseline from 4 samples from 3 cultures: N = 2057, CT 10min from 3 samples from 2 cultures: N = 642. P301L Baseline from 5 samples from 3 cultures: N = 2271, P301L 10min from 5 samples from 3 cultures: N = 1282. (G) – (L) Baseline from 6 samples and 4 cultures: N > 2000 synapses; 10 min from 5 samples and 3 cultures: N > 1200. **Statistics:** (C) Means from samples (each sample > 200 SVs) compared using a 2-sample t-test. (D)(E) 2-sampled t-test, (G)-(L) Mann-Whitney-U test. * = P< 0.05; ** = P < 0.01; *** = P< 0.001. Data points represent average values of binned data. Error bars represent standard error of binned average.

Here, we sought to determine if P301L neurons exhibit the same dynamic recycling pool changes during LTP and whether those changes follow the mechanistic changes hypothesized by the entropic force framework. We co-cultured primary hippocampal neurons and astrocytes from P301L mice (**Fig. 5A,B**, and **See Methods**) and used the same *θ*-burst protocol to induce plasticity in P301L cultures (**Fig. 1B)**. We also confirmed that the single SV pHluorin-VGlut1 intensity exhibits an increase (∼1.3 +/- 0.1) in P301L cultures compared to controls measured at the same time (**Fig. 5C** and **See Methods**), which is consistent with previously published results showing an increase in VGlut1 per SV in the presence of P301L tau.^52,55^

We followed the same pHluorin-VGlut1 protocol to image and quantify synaptic function (**Fig. 3B**). We then compared changes in synaptic function during the same early time period after plasticity induction (10 min). We first observed a significant reduction in VGlut1-pHluorin intensity during NH_4_Cl exposure at 10 min after induction (Blue Circles, **Fig. 5D**) compared to baseline (Black Squares, **Fig. 5D**). The intensity during 40 Hz stimulation also decreased at 10 min after induction (Blue Circles, **Fig. 5D**). These results suggest that the total number of SVs and the recycling pool density decrease after induction, as compared to the increase observed in normal neuronal cultures (**Fig. 3C**).

We then measured network-level averaged changes in SV number and spontaneous release rate for P301L cultures compared to controls. We observed a significant reduction in the total number of relative SVs in P301L cultures at 10 min compared to baseline (Red, **Fig. 5E**) and compared to control cultures (Black, **Fig. 5E**). This corresponded with the same changes observed in average spontaneous release (**Fig. 5F**). While these network level measurements have been the standard approach to understanding P301L induced changes but they do not indicate the underlying mechanisms leading to those changes.

To establish if P301L SV dynamics alter spontaneous release following the entropic force framework, we used the same analysis approach for control cultures (**Fig. 3**). First, we observed a slight but significant reduction in spontaneous release rate as a function of total synaptic vesicle number after induction (Blue Circles, **Fig. 5G**) as baseline (Black squares, **Fig. 5G**), which means the P301L neurons did ***not*** exhibit the same dynamic changes in spontaneous release observed in normal hippocampal cultures (**Fig. 3E**). The spontaneous release rate slightly decreased (p = 0.057) relative to stimulated release events at 10 min after stimulation (Blue Circles, **Fig. 5H**) relative to baseline (Black squares, **Fig. 5H**). Entropic force theory hypothesizes that the combined raw intensity results and stimulated/spontaneous release results suggest that P301L presynapses are losing SVs from the recycling pool.

To test whether recycling pool dynamics mediate the observed changes in spontaneous release behavior, we used the 40 Hz stimulated results to determine whether recycling pool size and dynamics change during plasticity. We observed that the fraction of SVs in the recycling pool decreased significantly as a function of the total number of SVs 10 min (Blue Circles, **Fig. 5J**) relative to baseline (Black squares, **Fig. 5J**). However, the spontaneous release rate remains the same as a function of recycling pool size for 10 min after induction (Blue Circles, **Fig. 5K**) relative to baseline (Black squares, **Fig. 5K**). We last compared changes in active zone mechanics to determine whether any functional/structural changes occur during plasticity. We observed that the release site number remained unchanged across recycling pool sizes (**Fig. 5L**). Further, the spontaneous release rate per release site is slightly increased at 10 min after induction (Blue Circles, **Fig. 5I**) relative to baseline (Black squares, **Fig. 5I**).

Taken together, these results show that, unlike normal neurons, P301L-expressing neurons exhibit a dynamic ***loss of SVs in the recycling pool during plasticity*** (**Fig. 5J**). This loss of recycling SVs competes with the increase in SV density during plasticity, leading to a constant relative spontaneous release rate with total SV number (**Fig. 5G**). Further, because the total size of the synapse is decreasing, this competition between loss of the recycling pool and changes in dynamics leads to a network-level decrease in spontaneous release rates during plasticity, which can be predicted by the entropic force framework.

We also highlight that the observed reduction in recycling pool SV fraction during plasticity represents a control test for the entropic force model. The reduction in spontaneous release rate (**Fig. 5G**) that corresponds with the reduction in SV density (**Fig. 5J**) still follows the entropic force framework even though it is the opposite behavior that occurs in control neurons (**Fig. 3E**). These combined results support the value of entropic force framework.

## Discussion

In this study, we show that the presynaptic spontaneous release rate is dynamically modulated during synaptic plasticity via changes in SV pool density, and the dynamic changes follow the entropic force model. We connected the dynamically changing presynaptic SV number with the inter-synaptic vesicle exchange (ISVE) pathway to show how presynapses can up/down regulate their SV pools (**Supplementary Material, Fig. S3**). Importantly, we showed how the dynamically changing spontaneous release rate is altered in the presence of the P301L tau mutation and may represent an early-stage pathway to neurodegeneration. These results both support previous observations of changing SV distribution^33^ and total SV number^62^ during plasticity, as well as provide significant new insight into the mechanics and consequences of these changes.

### Mechanism of Dynamically Changing SV Number

It has been well established that the actin cytoskeleton is the dominant structural mechanism that maintains SV number and turnover within the presynapse. Actin is essential for maintaining and regulating SV release during and after activity.^47,63,64^ Actin is important for SV recycling mechanics to maintain the recycling pool.^65,66^ We, along with many others, have shown that actin also maintains SV exchange between the presynapse and the surrounding axon.^46,67–74^ Thus, we hypothesize that actin dynamics during plasticity are the main mechanistic process by which SV number within presynapses changes dynamically. This hypothesis is consistent with, and ties together into a single coherent pathway, all previous observations on the role of actin before, during, and after activity. Further, this hypothesis also explains previous observations of up- or down-regulated SV pool sizes during LTP/LTD. There is a growing body of evidence showing changes in SV number and distribution during plasticity, and future studies should explore the specific role of actin structure/dynamics in regulating these dynamic changes.

### Meaning of Dynamically Changing SV Number

The role of dynamically changing SV numbers and pool distributions provides new insights into a combination of studies on synaptic plasticity. For example, previous studies have shown that the spontaneous release rate dynamically changes,^31^ total SV number changes,^32,33^ SV protein turnover rate changes,^75^ the overall size of pre/post synapse structure,^76^ and more recently, that the actin network dynamically restructures due to activity.^64^ However, these studies, which utilize post-synaptic measures (such as electrophysiology or PSD-95),^19,21,30^ or measure limited aspects of presynaptic changes using advanced fluorescent microscopy techniques.^27,77^ Our combined approaches in this study provide a mechanistic pathway by which all of these changes coordinate together to mediate presynaptic changes during plasticity: (i) the actin network restructures to more tightly bind SVs in the reserve pool; (ii) the increase in actin leads to more ISVE SVs to leave the axon and enter the presynapse increasing the density of SVs in the recycling pool; (iii) the increase in SV density leads to an increase in spontaneous release rates; (iv) the increase in spontaneous release rate leads to more SV recycling and thus protein degradation; (v) the actin network slowly depolymerizes releasing SVs from both the reserve pool and the recycling pool (used in step (iv) and replacing them with newer SVs). This single, coherent mechanistic pathway provides a framework for understanding the complex set of presynaptic changes that occur during plasticity.

### Value and Significance of Entropic Force Framework

We have previously shown that the entropic force framework can explain the complex biological processes during homeostasis that lead to the observed spontaneous release rate. Here, we show that the same theoretical framework can both explain and predict changes during plasticity. This approach significantly advances our understanding of plasticity because the location of presynaptic release sites has been shown to align with postsynaptic receptor sites.^2,22^ Further, spontaneous release has been shown to correlate with the strength of plastic changes that occur.^18,19,21^ The results in this study show that presynaptic changes in spontaneous release rate during plasticity can be modeled using entropic force theory. This suggests that the strength of postsynaptic changes, and thus changes in synaptic weight, can in turn be modeled more effectively. Future studies should explore how the temporal changes observed in this study relate to spatial changes in spontaneous release locations and, in turn, changes in postsynaptic receptor response.

### Consequences of P301L During Plasticity

The dynamic changes observed in normal neurons are inhibited during plasticity in the presence of P301L tau. This is consistent with previously observed changes in synaptic plasticity using ePhys approaches,^48,50,59^ as well as reductions in synaptic transmission in later stages of neurodegeneration.^48^ However, the results in this study provide a significant advance in our understanding of the mechanistic pathway by which reductions in synaptic function during plasticity occur during neurodegeneration. Our results suggest: (i) the mechanism of dynamically modulating SV number/density are inhibited in the presence of P301L tau; (ii) leading to a reduction in SV turnover during plasticity; (iii) reducing SV exocytosis/endocytosis efficiency; (iv) reducing synaptic function. The entropic force model provides a coherent theoretical framework for understanding and predicting how this process occurs. These results now provide specific molecular/mechanistic targets for future studies to explore to arrest or reverse the degenerative process.

### Limitations of the Present Study

While the present study provides a significant advancement in our understanding of presynaptic changes during plasticity, several limitations should be addressed in future studies. First, we used established hippocampal culture approaches that provide high spatial/temporal resolution but are limited because they do not replicate the exact circuit connections that occur physiologically. Second, the field stimulation approach we use in this study can induce a more dramatic change in SV dynamics and turnover than a more localized physiological response would induce. We note, however, that while our observed changes are likely amplified compared to physiological changes, we propose the same mechanisms still occur in physiologically relevant contexts, simply to a lesser degree. Third, the P301L tau results may be influenced by the known hyperexcitability process that occurs during the early stages of neurodegeneration.^50,58,59,78,79^ Combined, these limitations can lead to a more dramatic response in cell cultures than would occur under physiological conditions. However, these limitations do not alter the underlying mechanisms or theoretical entropic force framework. Future studies should explore these same processes under physiological conditions and/or under different types of plasticity induction.

## Acknowledgements

The authors would like to acknowledge funding for this study from Auburn University internal grant program. We acknowledge the assistance of Dr. Sanja Sviben and Greg Strout at the Washington University Center for Cellular Imaging (WUCCI) in EM studies, which is supported by Washington University School of Medicine, The Children’s Discovery Institute of Washington University and St. Louis Children’s Hospital (CDI-CORE-2015-505 and CDI-CORE-2019-813) and the Foundation for Barnes-Jewish Hospital (3770 and 4642). We would like to thank Washington University Hope Center Viral Vector Core for the production of the VGlut1-pHluorin vector used in this study. A subset of the control data was originally published in Cell Rep. 2025. Finally, the authors would like to acknowledge that *no part* of this study design, implementation, code development, data analysis, or figure generation involved the use of artificial intelligence, and all work is the authors’ own. Editing of the manuscript involved the use of artificial intelligence to correct minor textual issues.

## Author Contributions

RC and MR maintained mouse colonies and prepared cell cultures from the mouse colonies. RC performed cell culture preparations. PW, HS, and MM performed single SGC5 experiments, and MWG performed VGlut1-pHluorin and La-SEM experiments. PW and BP analyzed the pHluorin-VGlut1 data and La-SEM data and were blind to conditions. MWG, MM, and HS analyzed the SGC5 data and were blind to conditions. MWG developed computational models of synaptic vesicle mobility and spontaneous release rates. PW and BP performed data analysis of the P301L pHluorin-VGlut1 measurements. MWG and MR designed and funded the study. All authors contributed to the writing and editing of the manuscript.

## Methods

### Entropic Force Spontaneous Release Model

The entropic force framework assumes that the spontaneous release frequency is dominated by recycling pool dynamics. Recycling pool vesicles generate an entropic force, via vesicle diffusion, resulting in pressure the vesicles exert on the walls of the presynapse similar to an ideal gas. Recycling pool vesicles exert force on the vesicles tethered at the active zone as well as each other resulting in tethered vesicles occasionally fusing with the plasma membrane leading to an exocytosis event. The resulting spontaneous release frequency is then mediated by the pressure exerted following:

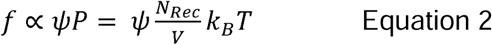

where N_rec_ is the number of mobile synaptic vesicles within the presynapse. The energy from each vesicle, which is what contributes to the force applied to the plasma membrane, is proportional to their speeds *k_B_T* ∝ *v*^2^. The number of SVs per volume is also quantified as the recycling pool density (*ρ_Rec_*). These combined metrics transform the equation into the form:

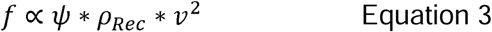

The **SPOntaneous Translation (SPOT) parameter** (ψ) represents the properties of the presynapse active zone release sites and local environment near the active zone (i.e. Ca^2+^ concentration) to translate the entropic force from the recycling pool into spontaneous release.

The active zone and release sites are essential in determining the probability of vesicle release.^2,80^ Increasing release sites will result in higher spontaneous release frequency. Here we assume that the contribution of SV force on spontaneous release is also a function of the number of release sites:

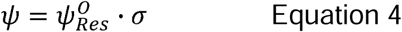

where *σ* is the number of release sites. Experimental results show that this is dependent upon the total vesicle number.

In general, presynapse size scales with the number of vesicles across different types of synapses. In the present study, we showed that presynapse volume increased linearly with the number of vesicles resulting in:

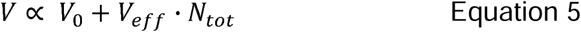

where V_0_ is the smallest possible presynapse volume and V_eff_ is the recycling pool volume increase per vesicle in the presynapse. Here the volume now represents the effective constrained volume of the Recycling pool, which we and others have shown is smaller than the total presynapse volume and constrained by the reserve pool.

We now turn to the origin of the SPOT parameter and its potential role in modeling changes in the active zone mechanics. This parameter can be thought of in a probabilistic framework, where each active zone molecular mechanism is represented as a probability:

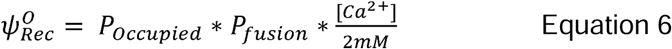

where the probability of a single SV tethered and primed at an individual release site is P_occupied_, and the probability of a primed SV can be fused with the plasma membrane in the presence of the entropic force is P_fusion_, and the relative Ca^2+^ concentration modulates the possibility of fusion. It is important to note that these probabilities are in units of time (i.e. probability per second).

### Entropic Force Theory Applied To Experimental Results

The entropic force model can be used to reproduce spontaneous release frequency measured as a function of stimulated release events by assuming relationships between the number of release sites (s) and observed stimulated release events (*N_stim events_*):

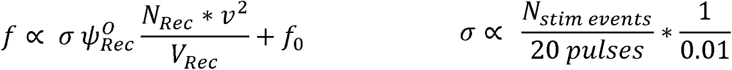

relationship between release sites and stimulated release events

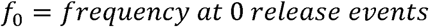

where 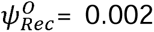, *N_Rec_* = 20, *V_Rec_* = 0.05, and *v*^2^ = (0.15 μm/sec)^2^, are taken from previously established values at baseline.^28^ This results in a linear relationship of spontaneous release frequency with the number of stimulated release events.

The entropic force model can also be used to reproduce spontaneous release frequency measured as a function of number of release sites directly by assuming relationships between the number of release sites (s) and number of recycling pool vesicles (*N_Rec_* ∼2.1 * *σ*, see fit from **Fig. 2**):

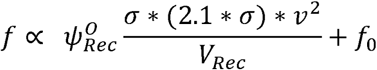

where this becomes a quadratic relationship with the number of release sites

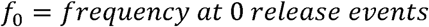

Where 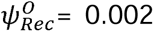, *V_Rec_* = 0.05, and *v*^2^ = (0.15 μm/sec)^2^. This results in a quadratic relationship of spontaneous release frequency with the number of release sites (*f* ∝ *σ*^2^).

Finally, the entropic force model can be used to reproduce spontaneous release frequency measured as a function of number of recycling pool vesicles directly by assuming relationships between the number of release sites (s) and number of recycling pool vesicles (*N_Rec_* ∼2.1 * *σ*):

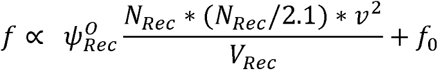

where this becomes a quadratic relationship with the number of release sites

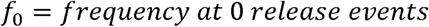

where 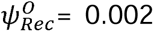, *V_Rec_* = 0.05, and *v*^2^ = (0.15 μm/sec)^2^. This results in a quadratic relationship of spontaneous release frequency with the recycling pool size (*f* ∝ *N_Rec_*^2^).

### Primary Rat/Mice Protocols

Timed pregnant rats were obtained from a commercial vendor (Sprague-Dawley, Charles River) and were held for at least 48 hours after arrival and before dissection. After 48 hours, the dam was euthanized using CO2 asphyxiation following NIH guidelines. Pup brains (at age E19) were then dissected, and the hippocampal portion of each lobe was removed. Pups were assumed to be evenly split between genders.

**Table 1:** Animal model used in each figure.

| Figure | Model Used |
| --- | --- |
| 1 | mouse |
| 2 | N/A |
| 3 | mouse |
| 4 | rat |
| 5 | P301L mouse |

Mice were created by crossing B6.Cg-Tg(Camk2a- tTA)1Mmay/DboJ (Stock No. <u>007004</u>) and FVB-*Fgf14^Tg(tetO-MAPT*P301L)4510Kha^* (Stock No. <u>015815</u>). All offspring were genotyped before postnatal day 4 (PND 4). Field ^44,45^ and gene-expressing mice were cultured separately, but at the same time as transgene-negative mice, so that changes in presynaptic function could be compared with transgenic-negative littermates. All mice were provided food and water ad libitum and housed under 12:12-hour light-dark cycles in humidity and temperature-controlled rooms. Mice used for dissection were assumed to be of mixed gender, a limitation of the present study.

We have previously used both the rats and mice species in the present study for the same experimental approaches,^43,44,81^ in order to support cross-species validation. The Auburn University Animal Care and Use Committee approved all experiments (protocol numbers: 2020-3715, 2020-3657, and 2020-3742). All protocols complied with the guidelines published in the NIH Guide for the Care and Use of Laboratory Animals.

### Cell Culture

Primary hippocampal neurons were plated on previously prepared astrocyte monolayer on prepared glass coverslips in Neuronal Growth Media consisting of 84% Minimum Essential Medium (Thermo Fisher), 9.6% Donor Equine Serum (Hyclone), 2% 1M Glucose in MEM (Thermo Fisher), 0.5% Penicillin/Streptomycin (Thermo Fisher), 1% N2 supplement (Thermo Fisher), 1% Sodium Pyruvate (Thermo Fisher), 2% 1M HEPES, by volume. 24 hours post dissection, the Neuronal Growth Media was replaced with Neurobasal media containing 96% Neurobasal-A Medium (Thermo Fisher), 2.5% B-27 supplement (Thermo Fisher), 0.3% Glutemax-1 (Thermo Fisher), 1.2% Penicillin/Streptomycin (Thermo Fisher), by volume. Plated cells were kept in an incubator until imaging (Div 14-21), with an additional 0.5 ml of Neurobasal media added every 7 days.

### Transduction Protocol

The VGlut1-pHluorin virus was obtained from the Washington University in St. Louis School of Medicine Hope Center Viral Vector Core. All cell cultures transduced with vGlut1-pHluorin at 3 DIV by replacing neurobasal medium with fresh 1 mL neurobasal medium containing pHluorin-VGlut1 viral titer. After 48 hours, the transduction medium was removed and replaced with fresh Neurobasal media not containing virus. Every 7 days another 0.5 ml of neurobasal media was removed and 0.5mL fresh neurobasal was added.

### Fluorescence Microscopy Experimental Approach

#### Microscope

Samples were imaged on a custom-built microscope with Ti2 base (Nikon), a 100x oil objective, Orca-Flash 4 CMOS camera with 6.5 x 6.5 *μ*m pixel size (Hamamatsu), and illuminated with a Sola LED light source. Excitation and Emission light were filtered using commercial cubes (GFP, Nikon). All experiments were performed with samples at physiological temperature 37°C within a whole-microscope incubator (OKO Labs) at DIV14–19.

#### SGC5 and pHluorin-VGlut1 media

Experiments were at physiological temperature 37°C and between DIV14–21. Transducted cultures were perfused with bath solution (140 mM NaCl, 2.5 mM KCl, 2 mM CaCl2, 4 mM MgCl2, 10 mM HEPES, 2 mM Glucose, 50 mM DL-AP5, 10 mM CNQX, pH adjusted to pH 7.4).^43,44,82–84^ SGC5 cultures involved a 4 min perfusion of media (140 mM NaCl, 2.5 mM KCl, 2 mM CaCl2, 4 mM MgCl2, 10 mM HEPES, 2 mM Glucose, 10 mM CNQX, pH adjusted to pH 7.25), after dye loading.^81,85^ All solutions were heated using a temperature controller attached to a multi-line solution heater (Warner Instruments). After induction but before imaging, cells were perfused with a bath solution (3, 8, 15 min) that did not contain post-synaptic blockers (DL-AP5 not CNQX) followed by 2 min perfusion with media containing post-synaptic blockers (DL-AP5 and CNQX). Ammonium Chloride solutions contained the above solution mixture plus 50 mM NH_4_Cl and pH-balanced to 7.4.

Stimulation and *θ*-Burst Protocol: Samples were electrically stimulated using parallel platinum electrodes with 1 msec pulse durations between 10-20 mA (or 5-10 V/cm) to follow previous established standards.^81,84–86^ Single SV SGC5 labeling involved a paired-pulse stimulation (5 msec between pulses) followed by a 30 sec delay. Bulk SV SGC5 labeling involved 200 pulses at 20 Hz followed by 1 min delay for dye loading. Plasticity induction via q-burst (**Fig. 1 B**) involved a bout of 5 pulses at 100 Hz followed by a delay of 20 msec repeated 5 times. This train was then repeated 3 times with 10 sec between each bout.

### Spontaneous and Stimulated Release Identification from pHluorin-VGlut1

Each presynaptic integrated intensity was run through a Kenedy-Chung filtering algorithm following established published parameters for VGlut1-pHluorin experiments.^83^ Background intensities were then fit using a normal distribution to determine the first standard deviation of noise. Release events were then identified as any change in intensity greater than one standard deviation of the background. The intensity for release event was calculated as the peak intensity minus the 3 frame average background intensity prior to the release event. The total time of intensity observed was calculated as the time from beginning of rising intensity to the time the intensity decreased below one standard deviation of the noise.

### Single SV Data Analysis

Single SV identification both the single and bulk raw tiff files were performed in imageJ and identified as bright puncta in the single movie that also had a larger corresponding bright puncta in the bulk movie (defined as a bulk intensity puncta >2 of the single movie puncta intensity). The position for each single SV and presynapse were recorded. Single SVs were secondarily confirmed using established computational track identification algorithms.^81,85,87,88^ High-resolution SV track positions were then run through our previously established correlation analysis algorithm to identify bouts of Fast, Diffusive, and Pausing behavior.^81,85^ The SV speed was determined as the average speed per frame averaged across all frames in a track. The average speed per condition is then calculated as the average speed across all tracks. The displacement was calculated as the average mean-square-displacement between 400 – 700 frames for each track calculated from the track position in the first frame. The displacement for each condition is then calculated as the average of all tracks measured for each condition.

### Large Area Scanning Electron Microscopy

Cells are prepared for La-SEM following our previously established protocol.^28,38^ Briefly, cultures were aldehyde fixed for 15 minutes at 37°C in modified Karnovsky’s fixative (2.5% glutaraldehyde, 2% paraformaldehyde, and cacodylate buffer) and stored in fixative at 4 °C overnight. Samples were washed with cacodylate buffer, followed by heavy metal incubation in a solution of 1.5% potassium ferrocyanide, 1% osmium tetroxide, and cacodylate buffer for 1 hour. Thiocarbohydrazide-osmium liganding (OTO) was performed, followed by a serial wash and incubation in 2% osmium tetroxide (aq). Samples were washed and incubated in 1% uranyl acetate (aq) overnight at 4°C. Samples were washed, and contrast was enhanced with en bloc staining in 20 mM lead aspartate at 60 °C. Samples then underwent a stepwise dehydration series in ice-cold ethanol for 10 minutes each (Ethanol/ddH20: 20%, 50%, 70%, 90%, 100%, and 100%). After dehydration, samples underwent serial resin infiltration at 2-hour intervals (Durcupan/ethanol: 25%, 50%, and 75%). Samples were then placed in 100% Durcupan resin overnight and then into fresh 100% Durcupan resin with the accelerator. Finally, samples were cured in a 60 °C oven for 48 hours.

Post resin curing, samples were sent to the Washington University Center for Cellular Imaging (WUCCI) where 50 nm thick five serial sections were cut in the cell culture growing plane and placed onto a silicon wafer chips. Large (∼ 300 x 300 µm) regions were imaged at 5 nm resolution in a FE-SEM (Zeiss Merlin, Oberkochen, Germany) using the ATLAS (Fibics, Ottowa, Canada) scan engine with a 5 µs dwell time and line average of 2. The SEM was operated at 8 KeV and 900 pA using the solid-state backscatter detector.

#### La-SEM Analysis

We utilized the same counting procedure we previously developed.^28^ Individual (300×300 *μ*m) SEM images were loaded into ImageJ and filtered using ImageJ’s Gaussian Blur filter with a rolling ball radius of 1. Presynapses were identified as intact regions with a grouping of SVs (>10). The area of each presynapse was found based on ellipticity calculations and by tracing the long and short axis of each presynapse. The active zone was identified as the darkest region along the perimeter where there were SVs within 50 nm. A *radial axis* was defined from the center of the active zone and orthogonal to the membrane (radial axis). Lines were drawn at 50 nm intervals perpendicular to the radial axis. The total number of SVs at each radial interval were recorded. The density of SVs at each 50 nm interval was calculated. The SVs defined as RRP are the number of SVs at 50 nm from the active zone. The Recycling pool SVs were defined as the SVs within the first 200 nm of the active zone.

### Fitting 40 Hz pHluorin-VGlut1 Intensity

We used the established multinomial model of presynaptic transmission^6,7,89^ to computationally model and fit the 40Hz data portion of the pHluorin protocol.^44^ We build a database of different pHluorin-VGlut1 intensity profiles as I(t) = n*p*q*t*N; where p is the probability of release per vesicle to a single stimulus (obtained from experimental sample response), n is the number of active zone sites (Fit to data), q is the intensity per vesicle (obtained from experiments) and then decays with time due to endocytosis (fit to data) and N is the number of stimulus per time-frame (here in units of frames).

Each experimental 40 Hz intensity profile compared to every computational profile in the database. The quality of fit for each profile was determined by a chi-square comparison. The modeled profile was chosen with the lowest chi-square. The parameters from the modeled profile (number of release sites, total recycling pool, etc…) were then combined with each synapse spontaneous and stimulated release events.

### Computational Simulation Model of SV mobility

#### Presynapse structure

Simulated presynaptic vesicle pool mobility was performed using a lattice-based model approach in python.^28^ Each site is designated as equal to the size of a single SV. Each time-step in the simulation is designated as 300 msec. The active zone is designated at one side of the lattice. The number of n-lattice sites along that side is equal to the number of release sites. We modeled the reserve pool as SVs fixed at lattice sites (> 5 sites for controls, and >3 lattice sites for 10 min, **Fig. 1K**). Recycling pool SVs followed mobility rules that reproduced random diffusive motion with elastic collisions. If a SV in the recycling pool occupies an active zone site, then a spontaneous release event is registered using a Monte-Carlo threshold algorithm (p = 0.002 per hit). At the end of each simulation, the average time between recorded exocytosis events is recorded as a rate per min. After 100 simulations, the average spontaneous rate per minute is calculated along with SEM for the rate.

### Computational Simulation of Axonal SV Trafficking and Differential Capture Mechanics

We computationally simulated SV trafficking along axons with a regular distribution of presynapses using our previously established computational approaches in python^81,90^:

i. we defined an axon as a one-dimensional lattice (100 nm/lattice site) and simulation time-steps equal to 0.2 sec each. The axon had a fixed length of 310 lattice site. The soma was defined at lattice site x = 0, and the axon growth cone was defined as a reflecting boundary where all SVs reversed direction and continued travelling.
ii. we defined presynapse locations at fixed distances along the axon (30 lattice sites, 3 microns) and fixed length (3 lattice sites). Individual SV tracks originated from a presynapse site for up to 20,000 time-steps (∼5.5 hrs modeled simulation time), and each track was independent of the other tracks. We simulated 200 tracks per presynapse, with 300 presynapses for a total of 6000 unique tracks. If a SV reaches the soma, then the simulation is ended. If the SV reaches the axon end (x = 310) it reverses direction.
iii. Any SV that reaches a designated presynapse location has a probability of capture (0.75). SVs that are captured pause for a random amount of time drawn from an exponential distribution
iv. , we simulated an entire axon of presynapses and SVs using a bootstrapping approach to randomly draw a single SV from each presynapse at fixed times during simulation. We randomly chose a single SV from each presynapse at a fixed time interval (every 30 time-steps). The SV track was then added to the aggregate axonal lattice independent of the other SVs. SVs were added to the simulation for 200 time-steps resulting in a total of 60,000 SVs per simulation.
v. the capture probability and pause-time dynamically varied as outlined in the results section under 3 different models.

### Data Aggregation, Binning, and Error-Bar Calculations

<u>The La-SEM</u> data were binned and error-bars calculated depending upon the independent parameter. (1) Network level calculations (Fig. 1 F,H) represent the average of all data measured, and error-bars represent the standard-error of the mean. (2) Cumulative distribution (Fig. 1G) were binned in 10 SV bin sizes. (3) Data as a function of “Total SV Number” (Fig. 1I) were binned in 10 SV sizes and error-bars represent the standard-error of the mean for each bin.

<u>The pHluorin-VGlut1</u> data were binned and error-bars calculated depending upon the independent parameter. (1) Network level calculations (Fig. 2 C,D; Fig. 3 D-E; Fig. 4 C-F) represent the average of all data measured, and error-bars represent the standard-error of the mean. (2) Data as a function of “Total SV Number” (Fig. 2 E,H,J; Fig. 4 G,J) were binned in 10 SV sizes for <100 SVs and 20 SV sizes for > 100 SVs; all error bars are 1/N. (3) Data as a function of “Recycling Pool Size” (Fig. 2 I; Fig. 4 I) or “Number of Release Sites” were binned in units of 5; all error bars are 1/N. (4) Data as a function of “Stimulated Release Events” were binned in single stimulated events; all error bars are 1/N.

### Statistical Analysis

KS-tests were used for cumulative distribution. Mann-Whitney U tests were used for repeated measures tests on measurements that exhibited trends as a function of the independent variable. For SGC5 comparisons between conditions, pair-wise t-Tests of raw data sets were used to determine significance of conditions.

For pHluorin-VGlut1 and La-SEM data, a multi-stage statistical analysis approach was performed:

i. Pair-wise t-Tests were used for whole raw data-set comparisons to determine if there were any statistical significance. Data sets that did not exhibit statistical significance were not compared in more detail and were reported as not significantly different.
ii. Data that exhibited statistical significance were then broken up into different ranges
iii. “Total SV Number” data (Fig. 2 E,H,J; Fig. 4 G,J) were compared for ranges <50 SVs and >50 SVs.
iv. SV Density data (Fig. 1H) were separated by Recycle/Reserve range
v. If data were statistically different for each range in (ii) then the Mann-Whitney U test was used to determine statistical difference reported for each range.
vi. If data were statistically different for all ranges (ii) then the Mann-Whitney U test was used and the total statistical difference from (i) was reported.

Statistical results were reported for each measurement for total data and/or data ranges where significances were observed in the figure legends. The number of animal litters, samples, and synapses used for statistical comparisons are listed in each figure legend for each condition.

## Supplementary Figures

### Section S1: Estimated Presynapse SV number as a function of Presynapse Size

**Figure S1:**
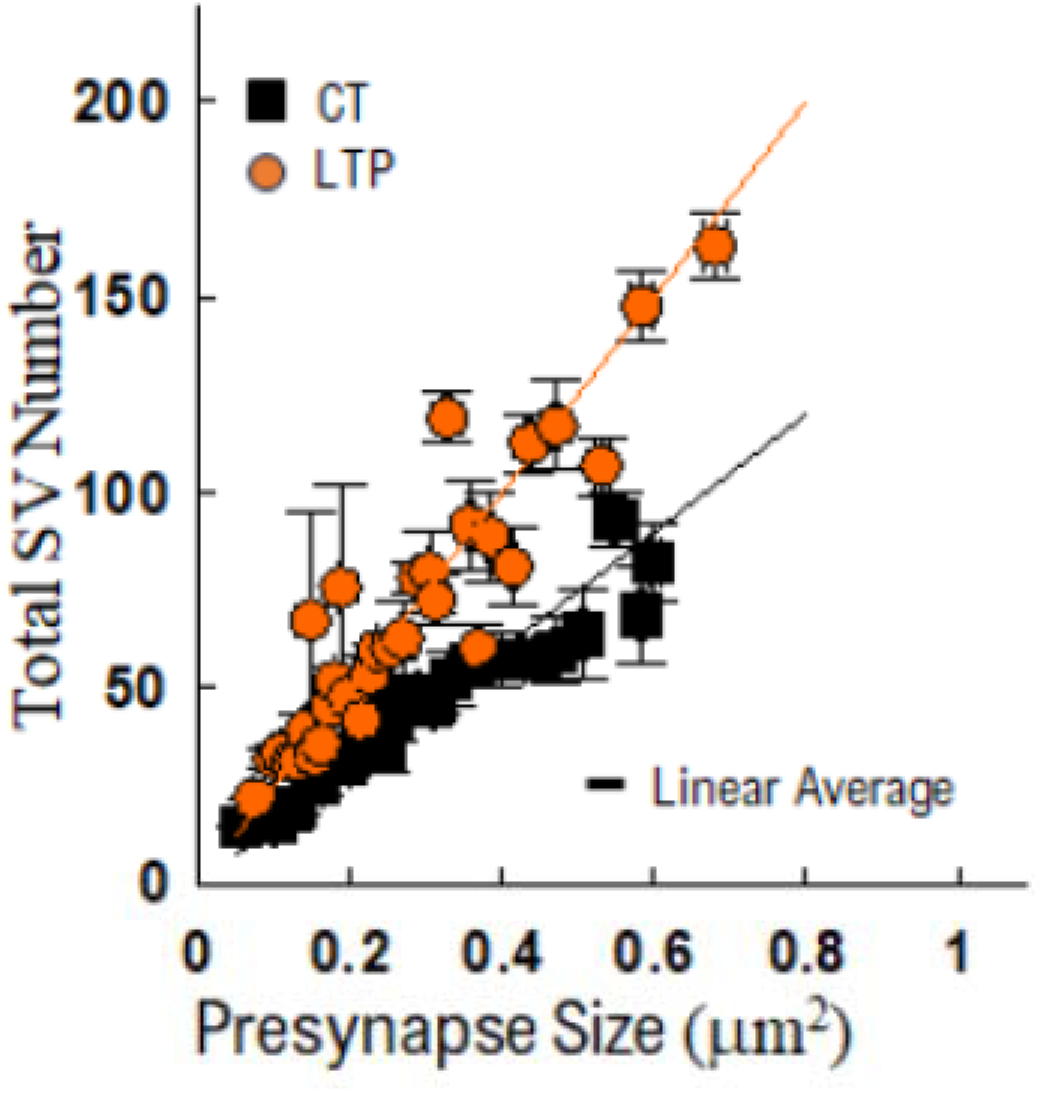
Total number of SVs within a presynapse as a function of presynapse size in SEM. The number of SVs and the geometric size of the synapse are calculated as independent parameters on a per-synapse basis. Data is binned as described in Methods. N_ct_ = 111 from two cultures; N_LTP_ = 165 from two cultures

In order to determine if the geometric size of the presynapse changes along with the number of SVs during plasticity, we compared the number of SVs as a function of size using SEM results. We observed an increase in the total number of SVs along with the size of the synapses at baseline (Black Squares, **Fig. S1**) that followed a linear relationship (Solid line, **Fig. S1**). Cultures exhibited the same relationship after plasticity induction (Orange Circles, **Fig. S1**); however, there was a scale increase in the relationship (Solid Line, **Fig. S1**). This increase means that the number of SVs per synapse is greater after induction for the same size presynapse. This result supports the hypothesis that the size of the presynapse does not significantly increase while the total number of SVs does increase, which leads to an increase in density (**Fig. 1H**).

### Section S2: Estimated release mechanics in P301L cultures

**Figure S2:**
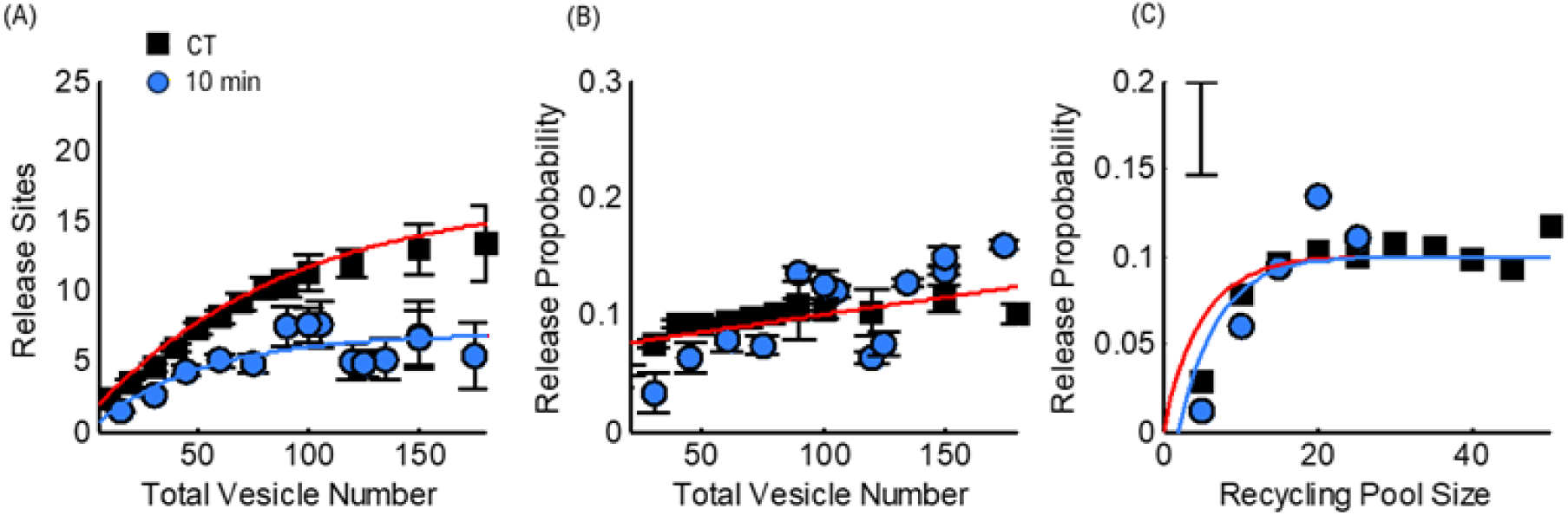
pHluolrin-VGlut2 data for P301L cultures at baseline and 10 min after induction. (A) The estimated number of release sites per total SV number for baseline (CT) and 10 min after induction (10 min) (B) The measured release probability per total SV number for baseline (CT) and 10 min after induction (10 min) (C) measured release probability per estimated recycling pool size for baseline (CT) and 10 min after induction (10 min) Baseline: N > 2000 synapses from 6 samples and 4 cultures; 10 min: N > 1200 from 5 samples and 3 cultures.

### Section S3: Synaptic Vesicle ISVE and Turnover Dynamically Change During Plasticity

**Figure S3:**
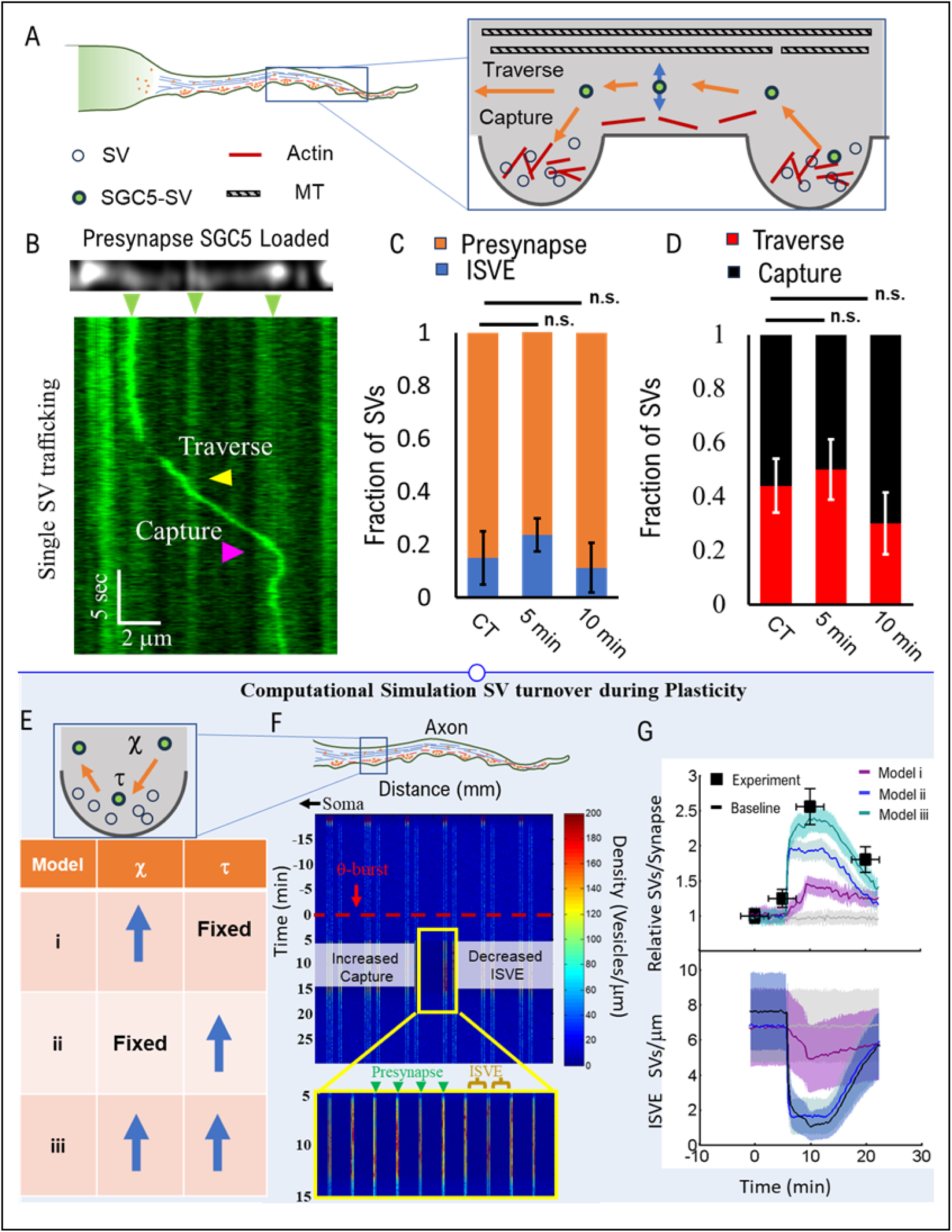
Single Labeled ISVE SV mobility Supports Dynamically Changing Total SV Number. (A) Cartoon Depiction of the labeled SGC5 SVs imaging approach during ISVE. Recently recycled SVs occasionally leave presynapses to exchange with neighboring presynapses or be trafficked back to the soma for degradation. (B) Example of a single SGC5 labeled SV in hippocampal cultures traversing (yellow arrow) or captured by (Magenta Arrow) presynapses with the majority of their SVs labeled (Green Arrows). (**C**) The fraction of identified ISVE SVs (Blue) as a function of total SVs observed (including SVs observed in Fig. 4) is typically less than 20%. The fraction does not change significantly over time after induction. (**D**) The fraction of ISVE SVs that are captured (Red), identified presynaptic locations slightly increase after activity relative to SVs that traverse (Black). (**E**) Example density plot of simulated SVs along an axon as a function of time. A simulated axonal bundle of presynapses and trafficking SVs has a steady-state number of SVs within presynapses and ISVE. We simulate a q-burst equivalent to experiments that yield dynamically changing ISVE capture/traverse rates and presynaptic dwell times. The simulations predict a significant reduction in ISVE SVs (lower density in the inset) and a significant increase in presynaptic SVs (increased SV density at pre-defined synapse locations). (**F**). (**G**) Comparison of simulated SVs per presynapse compared to experimentally measured pHluorin-VGlut1 results (Fig. 3D). (D) CT: N_SVs_ = 104, from 7 samples; 5 min: N_SVs_ = 61, from 6 samples; 10 min: N_SVs_ = 196, from 6 samples; (E) CT: N_events_ = 25, from 7 samples; 5 min: N_SVs_ = 56, from 6 samples; 10 min: N_SVs_ = 20, from 6 samples. Statistics: (C)(D) Test of Proportions. * = P< 0.05; ** = P < 0.01; *** = P< 0.001. Data points represent average values from aggregated SVs of multiple samples. Error bars represent

To address the mechanics of dynamic SV number and mobility, we quantified SV mobility during ISVE. Briefly, SVs that recently underwent exocytosis will leave the presynapse at a steady rate and begin ISVE along the axon during homeostasis (**Fig. S3A**).^46^ They then traverse along the axon to neighboring presynapses where they are captured depending upon which cytoskeletal network they traverse (**Fig. S3A**). We,^81^ and others,^67,69,72^ have shown that ISVE is a mechanism to maintain SV number within presynapses. We have also shown that differential ISVE trafficking is a mechanism for regulating the turnover of older SVs.^81^ Thus, we hypothesize that ISVE is the mechanistic pathway by which SV number and mobility dynamically change after induction.

Consistent with established ISVE behavior, we observe single SVs leave presynapses after labeling, and either traverse or be captured by neighboring presynapses (**Fig. S3B**). SVs will leave a presynapse, traverse the axonal cytoskeleton, and either be captured by the neighboring presynapse or traverse on toward the next presynapse. Thus, this approach can compare changes in the mechanics of SV behavior just after the recycling process as a function of time.

We tested the hypothesis that ISVE is the process by which presynapses dynamically change their pool size by first quantifying the fraction of ISVE SVs relative to the total number of single SVs observed. The fraction of ISVE SVs remains relatively constant (∼10-15%, **Fig. S3C**) initially after induction. Further, the fraction of ISVE SVs that were captured by presynapses during axonal motion increased 10 min after induction (**Fig. S3D**), which suggests that SVs were kept within presynapses more often, leading to an increase in total SV number per presynapse.

To understand how the ISVE metrics correlate with the experimentally observed dynamically changing SV number (**Fig. 1G and 3D**), we used our previously developed computational model of ISVE.^81^ Briefly, (i) SVs exit presynapses at a modeled rate, (ii) traverse the neuronal axon, (iii) are either captured (*χ*) or traverse (1-*χ*) neighboring presynapses with set capture probability, and (iv) remain within presynapses for random time (*τ*) until they exit again. Here, we modify the previous model to include dynamically changing capture probability and pause-time within a presynapse in three different conditions (models i – iii, **Fig. S3E**). SVs begin within synapses and exit at a fixed rate at baseline resulting in an equilibrium of SVs within presynapses as well as along the axon (**Fig. S3F**). The capture rate and/or the pause-time within presynapses started at baseline values previously established,^81^ and then changed dynamically after induction (dashed red line, **Fig. S3F**). We then determined the density of SVs along the axonal bundle as a function of time (**Fig. S3F**). We observed a dynamically changing ISVE and presynaptic density after induction consistent with experimental observations (inset, **Fig. S3G**).

We then quantitatively compared the simulated Total SV number per presynapse with pHluorin-VGlut1 over time and observed that our model reproduced the experimental results (Top panel, **Fig. S3G**). We found that model iii (a slightly increasing capture rate and doubling pause-time) reproduced the observed dynamic change in SV number as a function of time. We also compared the density of SVs along the axon (ISVE SVs) during the same simulated time-course and observed a significant dynamical change that matched the increase in presynaptic SV number (Bottom panel **Fig. S3G**).

These combined SGC5 and computational results support the hypothesis that a dynamically changing total presynaptic SV number is mediated by ISVE mechanics.

### Resources

| *Reagent or Resource* | *Source* | *Identifiers* |
| --- | --- | --- |
| *Chemicals, peptides, and recombinant proteins* |  |  |
| *DL-AP5 Sodium salt* | *Tocris* | *Cat. No. 3693* |
| *CNQX disodium Salt* | *Tocris* | *Cat. No. 1045* |
| *HEPES* | *Sigma* | *Cat. No. H4034* |
| *D-(+)-Glucose* | *Sigma* | *Cat. No. G 7021* |
| *Calcium chloride dihydrate* | *Sigma* | *Cat. No. 223506* |
| *Magnesium chloride hexahydrate* | *Sigma* | *Cat. No. M9272* |
| *Minimum Essential Media (MEM) - No Phenol Red* | *thermofisher* | *Cat. No. 51200-038* |
| *Characterized Fetal Bovine Serum* | *hyclone* | *Cat. No. SH30071.03* |
| *Penicillin- Streptomycin (5,000 U/mL)* | *thermofisher* | *Cat. No. 15070063* |
| *N-2 Supplement (100X)* | *thermofisher* | *Cat. No. 17502048* |
| *Donor Equine Serum* | *hyclone* | *Cat. No. SH30074.03* |
| *Sodium Pyruvate (100 mM)* | *thermofisher* | *Cat. No. 11360-070* |
| *Neurobasal- A Medium* | *thermofisher* | *Cat. No. 10888–022* |
| *B-27 Supplement (50x), serum free* | *thermofisher* | *Cat. No. 17504–044* |
| *GlutaMAX supplement* | *thermofisher* | *Cat. No. 35050061* |
| *Earle's Balanced Salts* | *sigmaaldrich* | *Cat. No. E3024* |
| *Corning Collagen I, Rat* | *FISHER SCIENTIFIC* | *Cat. No. 354236* |
| *Cover Glass* | *FISHER SCIENTIFIC* | *Cat. No. 12-545-80* |
| *Papain* | *Worthington Biochemical* | *Cat. No. LS003126* |
| *PDL (poly-D-lysine)* | *BD Biosciences* | *Cat. No. 40210* |
| *Trypsin-EDTA (0.05%), phenol red* | *thermofisher* | *Cat. No. 25300–054* |
| *Nocodazole* | *sigmaaldrich* | *Cat. No. M1404* |
| *HEPES Buffer 1M solution* | *Corning* | *Cat. No. 25-060-CI* |
| *SGC5* | *Biotium* | *Cat. No. 70057* |
| *Latrunculin A* | *AbcamBiochemical* | *Cat. No. ab144290* |
| *Forskolin* | *Millipore Sigma* |  |
| *Experimental Models: Organism/strains* |  |  |
| *Sprague Dawley Timed Pregnant Dam* | *Charles River* | *Strain# 001* |
| *tTa mouse line: B6.Cg-Tg(Camk2a-tTA)1Mmay/DboJ* | *Jackson Labs* | *Stock No. 007004* |
| *FVB-Fgf14Tg(tetO-MAPT\*P301L)4510Kha* | *Jackson Labs* | *Stock No. 015815* |
| *3xTg-AD mouse line* | *Jackson Labs* | *Strain # 004807* |
| *Software and algorithms* |  |  |
| *MATLAB* | *MathWorks* | *RRID:SCR_001622* |
| *ImageJ* | *https://imagej.nih.gov/ij/* | *RRID:SCR_003070* |
| *Fiji* | *http://fiji.sc* | *RRID:SCR_002285* |
| *Recombinant DNA* |  |  |
| *VGluT1-pHluorin* | *Drs. Robert Edwards and Susan Voglmaier (UCSF)* | *Genetically encoded optical indicator of vesicle release and recycling* |

